# Synaptic density and relative connectivity conservation maintain circuit stability across development

**DOI:** 10.1101/2025.07.26.666968

**Authors:** Ingo Fritz, Feiyu Wang, Ricardo Chirif, Nikos Malakasis, Julijana Gjorgjieva, André Ferreira Castro

**Affiliations:** School of Life Sciences, Technical University of Munich, Freising, Germany

## Abstract

As bodies grow during postembryonic and postnatal development, nervous systems must expand to preserve circuit integrity. To investigate how circuits retain stable wiring and function throughout development, we combined synaptic-level resolution electron microscopy (EM) with computational modeling in the *Drosophila* larval nociceptive system. Based on EM data, we generated the “contactome”—the set of synaptic membrane contacts—of this circuit across development to evaluate how different mechanisms contribute to wiring stability. Specifically, we investigated three mechanisms: correlation-based plasticity and synaptic scaling, which modify synaptic strength, and structural plasticity, which preserves synaptic density. We found that synaptic sizes remain largely stable across development, and synapses between the same pre- and postsynaptic neurons do not correlate in size, suggesting that synaptic scaling and correlation-based plasticity play a limited role in shaping connectivity. In contrast, dendritic synaptic density remains invariant despite a previously reported fivefold increase in neuron size and synapse number. This conservation requires increased axonal presynaptic density to compensate for unequal axonal and dendritic growth. As neurons grow, this adjustment is necessary to maintain the relative synaptic input associated with each presynaptic partner across development. Our EM analysis and modeling show that conserving relative connectivity and synaptic density is sufficient to maintain consistent postsynaptic responses across development, highlighting these conserved structural features as key contributors to circuit stability during growth.

## Introduction

From nematodes to humans, developing nervous systems undergo extensive growth and differentiation (***McDole et al., 2018; Doe, 2017; Chhetri et al., 2015; La Manno et al., 2021***). During early development, neurons extend and retract neurites, navigate complex environments, identify appropriate partners, and form synapses with remarkable specificity (***Heckman and Doe, 2022; Shree et al., 2022; Gour et al., 2021; Valdes-Aleman et al., 2021; Harth et al., 2024; Udvary et al., 2022; Kirchner et al., 2025***). Although significant progress has been made in understanding how neural circuits are initially established, much less is known about the principles that preserve function as nervous systems grow postembryonically and postnatally (***Kaiser, 2017; Heckman and Doe, 2021***).

Despite dramatic anatomical changes, circuit function is often conserved as animals grow (***Bucher et al., 2005; Murphey and Chiba, 1990; Almeida-Carvalho et al., 2017***). This functional maintenance ultimately depends on the combination of neuronal morphology, synaptic connectivity, and ion channel properties (***Marder and Goaillard, 2006***). Although theoretical and experimental studies have shown that adaptations in intrinsic functional properties, such as ion channel expression and distribution, can provide functional stability during growth (***Gorur-Shandilya et al., 2020; Prinz et al., 2004; Mease et al., 2013; Gjorgjieva et al., 2014***), recent electron microscopy (EM) studies have highlighted the conservation of connectivity patterns as another mechanism for maintaining circuit function (***Gerhard et al., 2017; Witvliet et al., 2021***). EM reconstructions of model systems such as *C. elegans, Drosophila*, and mouse have shown that the proportion of synapses a neuron receives from a specific partner, relative to the total number of inputs it receives – hereafter named relative connectivity – is conserved across cell types (***Gour et al., 2021; Schneider-Mizell et al., 2025***), individuals within species (***Schlegel et al., 2024***), hemispheres (***Pedigo et al., 2023***), and even stages of development (***Gerhard et al., 2017; Witvliet et al., 2021***). This conservation of relative connectivity hints at developmental mechanisms that shape neuronal wiring and suggests that maintaining input proportions is essential to preserve circuit function throughout development. These observations raise two key questions: (1) How does relative connectivity contribute to functional stability? (2) If relative connectivity is conserved, through what structural or activity-dependent processes is synaptic strength or number regulated during neuronal growth?

Several mechanisms have been proposed to account for the structural and functional stability of neural circuits during development. Two of the most widely studied are correlation-based synaptic plasticity (***Hebb, 2005; Miller et al., 1989; Oja, 1982; Bliss and Lømo, 1973; Ueno et al., 2013; Ataman et al., 2008; Markram et al., 1997; Gjorgjieva et al., 2009***) and synaptic scaling (***Turrigiano, 2012; Keck et al., 2013; Turrigiano and Nelson, 2004; Müller and Davis, 2012; Baines et al., 2001; Davis, 2013***), both of which provide activity-dependent strategies for regulating synaptic strength as neurons grow. Structural plasticity mechanisms that preserve synaptic density during growth, have also been proposed to contribute to functional stability without requiring changes in synaptic strength (***Royer and Paré, 2003; Bourne and Harris, 2011; Nelson et al., 2024; Lu et al., 2025; Kirchner and Gjorgjieva, 2021; Kirchner et al., 2025***).

Correlation-based plasticity, such as Hebbian rate-based or spike-timing–dependent learning rules, have long been used to study experience-dependent synaptic refinement in the brain (***Sejnowski and Tesauro, 1989***). These mechanisms have been shown to shape connectivity through the selective potentiation and depression of synapses based on the activity patterns of pre- and postsynaptic neurons (***Abbott and Nelson, 2000; Malenka and Bear, 2004; Ueno et al., 2013; Ataman et al., 2008; Markram et al., 1997***). Synapses with shared activity history may converge in strength and size, a signature that has been observed in synaptic-resolution EM studies, particularly when synapses from the same presynaptic neuron target the same dendritic branch (***Bartol et al., 2015; Dorkenwald et al., 2022; Motta et al., 2019***). Given that synaptic size correlates with strength (***Holler et al., 2021; DiAntonio et al., 1999***), these findings suggest that traces of correlation-based plasticity can be identified through connectomics snapshot experiments. However, it remains unclear whether correlation-based plasticity alone can conserve relative input proportions as neurons grow (***DeBello, 2008; Chklovskii et al., 2004***).

Second, homeostatic mechanisms are thought to stabilize neural circuit output by adjusting synaptic strengths in response to changes in activity (***Tien and Kerschensteiner, 2018; Turrigiano et al., 1998***). One prominent form of homeostatic plasticity is synaptic scaling, which ensures that synapses are scaled up to maintain neuronal excitability within functional limits as circuits mature (***Turrigiano and Nelson, 2004; Müller and Davis, 2012; Baines et al., 2001; Davis, 2013***). As neurons grow, their increased membrane surface area leads to lower input resistance in postsynaptic partners, potentially requiring stronger synaptic currents to sustain excitation (***Sterling and Laughlin, 2015; Dayan and Abbott, 2002***). Synaptic upscaling may compensate for these changes by proportionally increasing synaptic sizes, leading to a rightward shift in synaptic size distributions during growth and maintaining excitability at a constant level (***Hobbiss et al., 2018; Lichtman et al., 1987; Davis and Goodman, 1998; Govind and Pearce, 1981***). In particular, because synaptic scaling adjusts synapses proportionally, it could conserve relative connectivity and maintain the shape of the distribution of synaptic sizes established early in development.

In an accompanying comparative connectomics study, we have shown that synaptic density, i.e., the number of synapses per unit dendritic length, remains remarkably constant across species and brain regions (***Ferreira Castro and Cardona, 2024***). Besides mechanisms that change synaptic strength, this property has been proposed as a strategy to stabilize neuronal excitability (***Cuntz et al., 2021***). Intuitively, as dendrites expand, they can accommodate more synapses, increasing total synaptic input. This can offset the reduction in input resistance that accompanies larger neuronal size, thereby maintaining postsynaptic excitability. Computational models suggest that when synaptic density is constant, neuronal voltage responses and firing rates scale with the relative input from each presynaptic partner, rather than with the absolute number of synapses in that connection (***Ferreira Castro and Cardona, 2024; Cuntz et al., 2021***). However, what mechanisms might preserve synaptic density during rapid neuronal expansion in postembryonic and postnatal development is unclear. In particular, whether this invariance can be maintained through proportional scaling of axons and dendrites or requires active structural plasticity remains an open question (***Lu et al., 2025; Tripodi et al., 2008; Udvary et al., 2022; Harth et al., 2024; Kirchner and Gjorgjieva, 2021; Kirchner et al., 2025***).

Addressing these questions requires mapping neural circuitry at synaptic-level resolution across development, quantifying synaptic sizes between identified partners, and linking structural changes to neuronal excitability and behavior. The *Drosophila* larval nociceptive circuit, functionally analogous to the mammalian pain pathway, serves as an ideal model due to its well-defined connectivity, anatomical tractability, and behavioral stability. This circuit plays a vital role in larval survival by initiating rapid escape behaviors, such as rolling away from predatory attacks (***Figure 1***A) (***Hwang et al., 2007; Ohyama et al., 2015; Jovanic et al., 2016***). These responses are mediated by six multidendritic arborization class IV (mdIV) neurons, which span the body wall segments and are dorsoventrally segregated, with three neurons per segment on each side (***Figure 1***B). These neurons detect nociceptive stimuli through their dendritic arbors and transmit signals via axonal projections to downstream neurons in the ventral nerve cord (VNC), where sensory integration occurs (***Jan and Jan, 2010***). Among the primary recipients of mdIV signals are local neurons (LNs), which predominantly process information within local circuits and exhibit preferential connectivity with specific mdIV neurons encoding distinct body regions (***Figure 1***B) (***Ohyama et al., 2015; Gerhard et al., 2017***). Despite a two-order-of-magnitude increase in body wall surface area between the first (L1) and last (L3) instar phases of postembryonic larval development (***Keshishian et al., 1993***), behavioral output remains stable, implying that neural circuits must adapt structurally to maintain function (***Almeida-Carvalho et al., 2017***). Recent EM reconstructions of this circuit across developmental stages have revealed that, despite a five-fold increase in neuron size and synapse number, the relative connectivity between specific partners remains stable (***Figure 1***C) (***Gerhard et al., 2017***). This combination of precise anatomical mapping and developmental stability makes the *Drosophila* nociceptive circuit a unique system for studying the principles that govern circuit maintenance during postembryonic growth.

**Figure 1.**
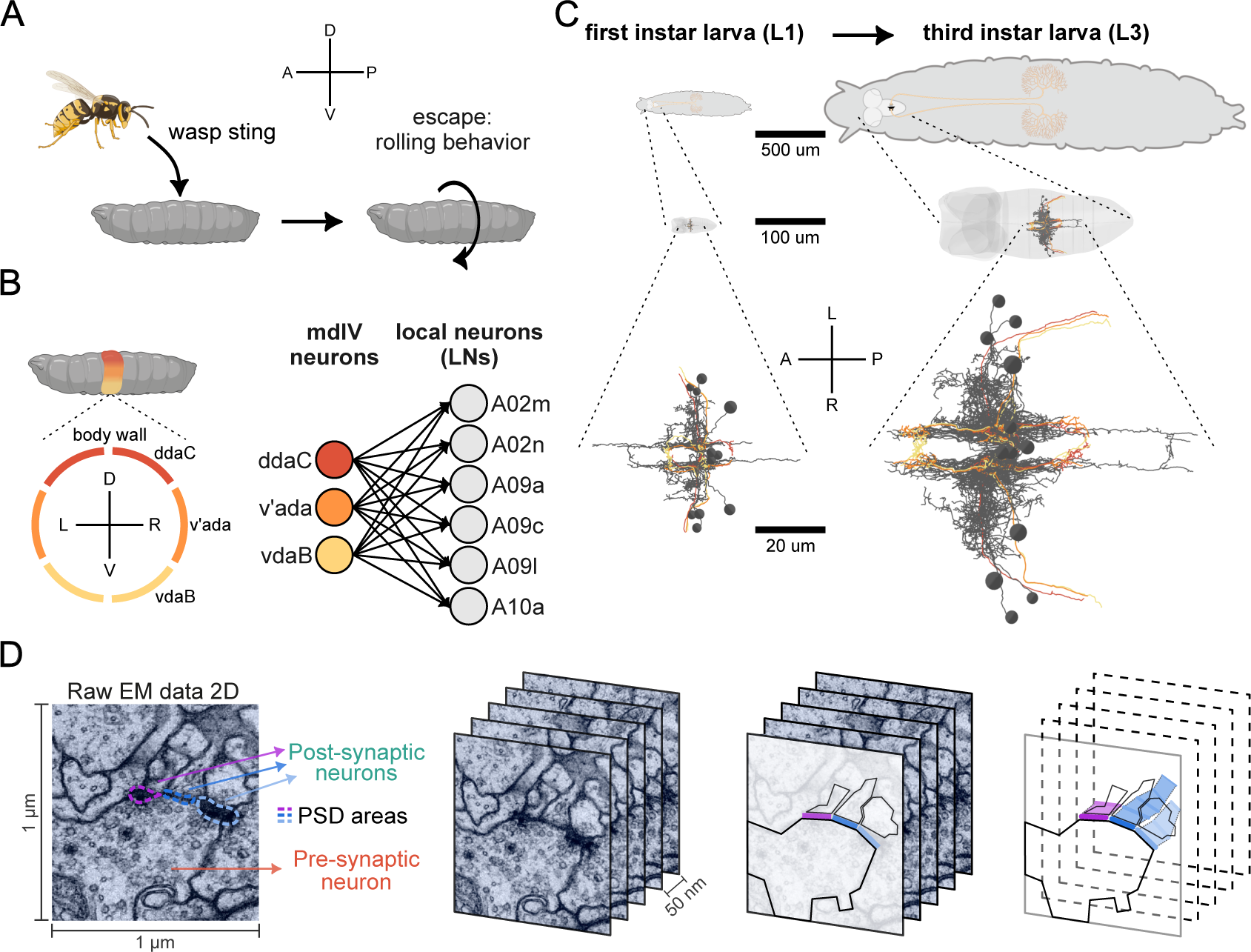
Connectome and contactome of the nociceptive circuit in *Drosophila* larva at developmental stages L1 and L3. (**A**) Illustration of the escape behavior triggered by nociceptive or damaging stimuli sensation. A-anterior; P-posterior; D-dorsal; V-ventral. (**B**) Circular arrangement of the nociceptive dendrites (ddac, v’ada and vdaB) and their projections to LNs (A02m, A02n, A09a, A09c, A09l and A10a). (**C**) Comparative visualization of the larva body (top), larva nervous system (middle) and nociceptive circuit (bottom) in the L1 and L3 developmental stages. A-anterior; P-posterior; L-left; R-right. (**D**) Postsynaptic density (PSD) segmentation pipeline. Left: dashed lines outline distinct PSD areas segmented from ssTEM images used in the analysis. Right: schematic showing cross-sectional views of a neurite, highlighting pre- and postsynaptic sites with color-coded representations of the neurite and its synaptic partners.

Despite these insights, it remains unclear how the nociceptive circuit maintains stable function across the dramatic growth of postembryonic development. To address this gap, we analyzed synaptic-level resolution EM datasets of the *Drosophila* larval nociceptive circuit across development to identify structural synaptic features that support circuit stability (***Gerhard et al., 2017***). To this end, we generated the contactome of the circuit, enabling us to map the size and distribution of synaptic contacts between identified neuron pairs. Surprisingly, across development, synaptic size distributions remain largely stable across postembryonic stages, and synaptic sizes between the same pre- and postsynaptic partners do not correlate. These findings suggest that mechanisms which only alter synaptic strength, such as synaptic upscaling and correlation-based plasticity, may not be the main drivers in shaping circuit stability. We also found that postsynaptic dendritic density remains stable across growth. Morphological analysis shows that this stability cannot be explained by spatial proximity alone, but requires an increased axonal presynaptic den-sity to maintain postsynaptic dendritic density and relative connectivity. Finally, EM-constrained single-cell computational modeling shows that these conserved features are sufficient to produce consistent postsynaptic functional responses, independent of developmental changes in neuronal size or morphology. Together, these results support a model in which synaptic density and relative connectivity may serve as structural substrates for developmental circuit stability.

## Results

To investigate circuit functional stability during postembryonic development, we quantified synaptic sizes across two connectomic datasets in first (L1) and third (L3) instar larvae. Given that synapses in the *Drosophila* nervous system are predominantly polyadic, that is, one pre-to many postsynapses, the area of the postsynaptic membrane is a reliable proxy of synaptic strength (***Barnes et al., 2022***). We measured postsynaptic densities (PSDs), protein-dense specializations at the postsynaptic membrane that appear as electron-dense regions in EM images, not to be confused with dendritic cable synaptic density. Measurements were done for all mdIV-to-LN connections (*N*_*L*1_ = 319; *N*_*L*3_ = 1562 synapses) in serial section transmission electron microscopy (ssTEM) image stacks from both L1 and L3 stages (***Figure 1***D; see Methods and Materials: Post synaptic density (PSD) segmentation and measurement). By segmenting PSD areas across multiple image slices (z-axis of the connectome), we estimated overall PSD area of each connection (***Figure 1***D) (***Barnes et al., 2022***).

### The distribution of synaptic sizes remains largely stable over development

We first evaluated how PSD areas, hereafter referred to as synaptic size, change over development and whether synaptic scaling compensates for dendritic growth by increasing them. If synaptic up-scaling were strongly engaged during development, we would expect a shift toward larger synaptic sizes, reflecting a multiplicative adjustment to offset the reduced input resistance of growing neurons (***Figure 2***A) (***Loewenstein et al., 2011***). Instead, we observed a significant reduction of approximately 19% in median PSD areas (***Figure 2***B, L1 vs. L3: *p* < 0.001).

**Figure 2.**
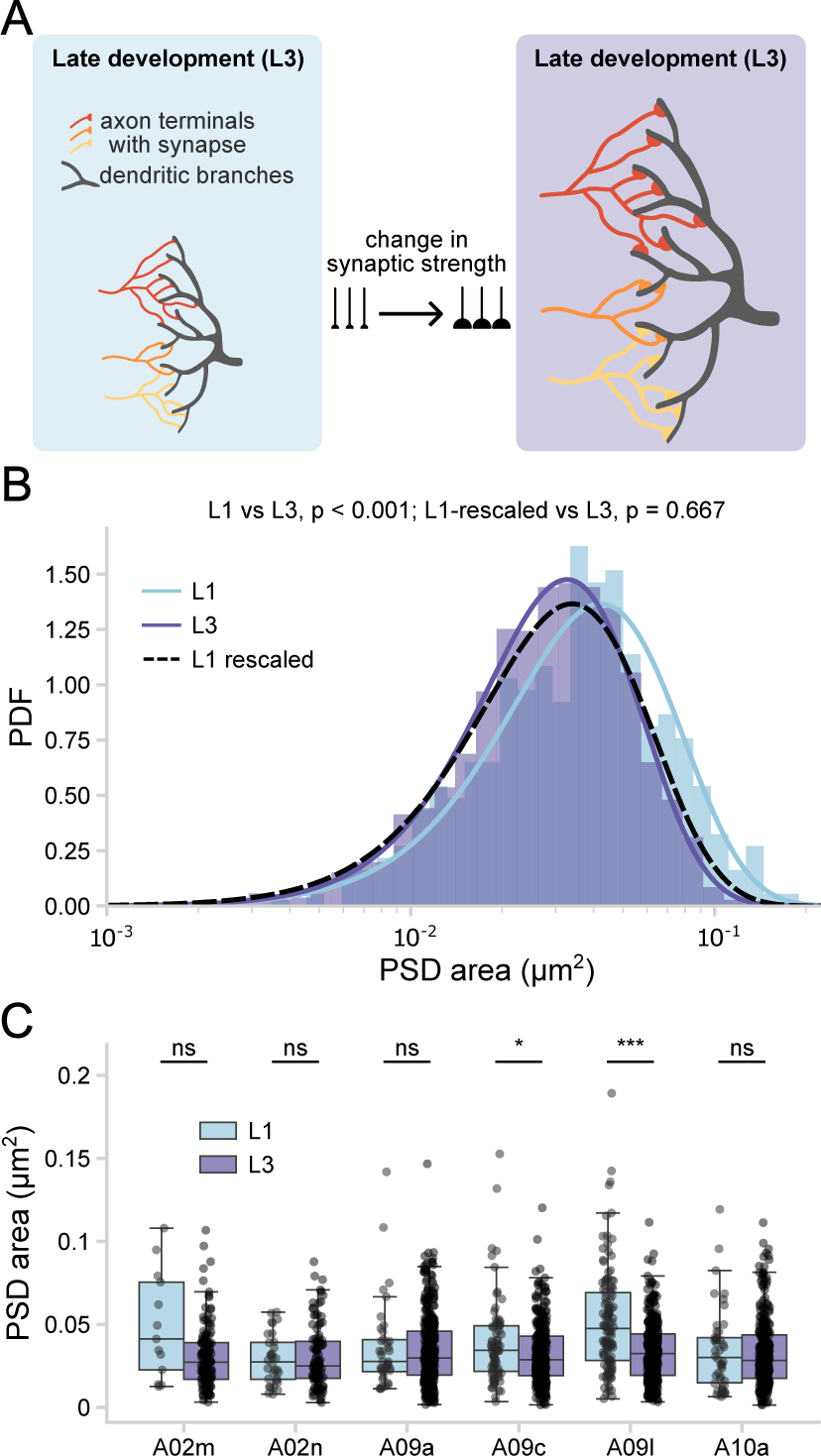
Connectomic mapping of synaptic size distributions across development. (**A**) Schematic illustration of expected shifts in synaptic sizes under homeostatic synaptic upscaling mechanisms for the L1 (blue) and L3 (purple) developmental stage. (**B**) Synaptic size (PSD area, log-scaled) distributions in L1 (N=319; blue; across all connections) and L3 (N=1562; purple; across all connections) datasets fitted with gamma curves. Dashed line shows the L1 distribution after applying a multiplicative rescaling factor (*s* = 0.81) to all PSD area values, aligning the L1 median to that of L3 while preserving the gamma shape. (**C**) Box plots of synaptic area distributions (data points) for individual LN subtypes in L1 (light blue) and L3 (purple). Synaptic size remains stable across most subtypes, with only A09c and A09l exhibiting significant differences.

Synaptic sizes were well fit by both gamma (KS statistic = 0.055, *p* = 0.281) and log-normal (KS statistic = 0.043, *p* = 0.584) models in L1, while the L3 distribution was best fit by a gamma model (KS statistic = 0.014, *p* = 0.920) and showed a significant deviation from log-normality (KS statistic = 0.053, *p* < 0.001). This shift suggests a narrowing of synaptic size variability over development (***Figure 2***B; ***Figure S1***). This is supported by a reduction of the coefficient of variation from L1 to L3 (L1 *CV* = 0.69 vs. L3 *CV* = 0.59: F statistic = 52.38, *p* < 0.001). Although not statistically significant, intra-individual differences in median PSD area were larger in L1 (≈ 11.5%, *p* = 0.29) than in L3 (≈ 0.5%, *p* = 0.65), consistent with a developmental reduction in variability.

To evaluate whether these differences reflect a change in scale, we applied a multiplicative scaling transformation to the L1 distribution (*s* = 0.81), uniformly scaling all PSD area values to match the median of L3. This transformation preserved the overall distribution shape (***Figure 2***B, scaled L1 vs. L3: KS statistic = 0.044, *p* = 0.667), suggesting that the median size difference and reduced variability could reflect a proportional change in size.

To further evaluate whether this shift was global or cell-type-specific, we compared synaptic size distributions across LN subtypes. Most subtypes remained stable between L1 and L3, with only two showing a significant reduction in median size (***Figure 2***C). This pattern argues against a global synaptic downscaling mechanism and instead suggests that the observed difference in median size may reflect either subtype-specific developmental changes or inter-individual variability, given that the L1 and L3 datasets were derived from different animals (***Gerhard et al., 2017***).

Together, these results suggest that synaptic sizes remain largely stable across development, with only modest changes in size and variability. The log-normal fits of synaptic sizes may reflect multiplicative dynamics of synaptic strength changes during development (***Loewenstein et al., 2011; Rößler et al., 2023; Piazza et al., 2025***), while gamma distributions of synaptic sizes can arise from additive or bounded synaptic strength changes (***Karbowski and Urban, 2023; Benavides-Piccione et al., 2013***). Although our data are consistent with synaptic downscaling, they do not support the upscaling that would be necessary to account for neuronal growth. While we cannot rule out a role for such mechanisms, they seem unlikely to be the dominant strategy for maintaining excitability and relative connectivity in this circuit (***Turrigiano and Nelson, 2004; Gjorgjieva et al., 2016***).

### Synapses sharing the pre- and postsynaptic neurons do not correlate in size

The observed shift in PSD area with neuronal growth toward smaller synapses is consistent with synaptic downscaling. Synaptic scaling is often thought to operate alongside correlation-based plasticity to stabilize connectivity and avoid runaway instability (***Zenke et al., 2017; Turrigiano***, ***2012; Wu et al., 2020***). This relationship raises the possibility that, even in the absence of up-scaling, correlation-based mechanisms, which change synaptic strength based on correlated pre- and postsynaptic activity, may still shape connectivity. To explore this possibility, we examined whether anatomical signatures of correlation-based plasticity are present in the mdIV-to-LN circuit—features that could emerge even under homeostatic conditions.

We first asked whether differences in synaptic size might simply reflect connection-specific properties rather than activity-dependent plasticity. To assess this, we examined whether pref-erential mdIV-to-LN connections, that is, those with higher synapse counts, might also have larger synapses (***Gerhard et al., 2017***). For each LN, we ranked its connected mdIV neurons based on the number of synapses they formed, assigning rank 1 to the strongest (most synapses) and higher ranks to progressively weaker (fewer synapses) connections (up to 5 mdIVs per LN; ***Figure 3***A; see Methods and Materials: Evaluating traces of plasticity on postsynaptic densities). We then compared mean PSD area across ranks to assess whether synaptic size varies systematically with connection rank throughout development. In both L1 and L3, the average number of synapses per connection decreases with increasing rank, confirming the intended ranking scheme (***Figure 3***B,C). However, mean PSD area per connection (black dots) shows no significant correlation with rank (L1: *r* = 0.01, *p* = 0.97; L3: *r* = −0.21, *p* = 0.13), suggesting that synapse size is not systematically modulated according to preferred connection at this level of organization.

**Figure 3.**
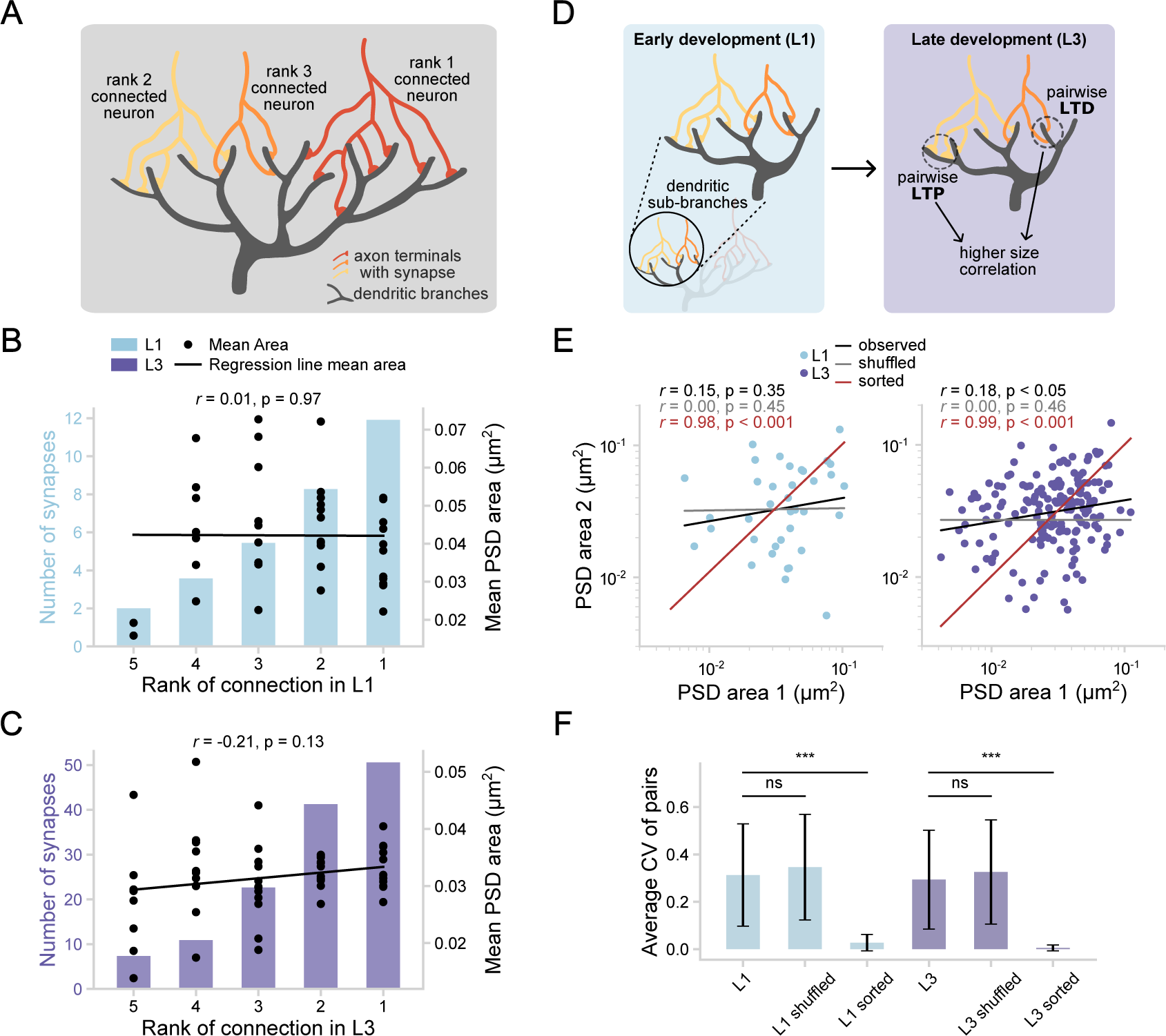
Evaluating traces of correlation-based plasticity across development. (**A**) Schematic illustrating the ranking of mdIV neurons connected to a single LN based on synapse count. The mdIV neuron forming the most synapses is assigned rank 1 (preferential connection), followed by those with progressively fewer synapses (ranks 2, 3, etc.). (**B, C**) Connectivity rank analysis of mdIV-to-LN synaps es in L1 (top, **B**) and L3 (bottom, **C**). Bars show the mean number of synapses per connection pooled across all LNs by ranking (left y-axis). Black dots represent the mean PSD area for each individual connection (right y-axis). Black line shows the regression of mean PSD area versus connection rank. (**D**) Schematic illustrating synaptic pairs originating from the same axon and targeting the same dendritic branch. Under correlation-based plasticity, such pairs are expected to exhibit greater size similarity. (**E**) Correlation of PSD areas from synaptic pairs within the same dendritic branch and sharing the same pre- and post-neuron in L1 (N=42; left) and L3 (N=172; right) datasets. Black line: linear fit of observed data; gray line: linear fit of shuffled control, with 1000 shuffled versions; red line: linear fit of sorted pairs (maximum correlation). (**F**) Mean coefficient of variation (CV ± SD) for PSD area pairs in observed data, shuffled, and sorted controls.

We next examined whether anatomical signatures consistent with Hebbian-like correlation-based plasticity are detectable at the LN dendritic branch-level. This is key because stochastic variability may confound quantification when pooling PSD areas across entire connections that span multiple postsynaptic dendritic branches (***Bartol et al., 2015***). Along these lines, studies in the mouse somatosensory cortex and rat hippocampal CA1 have shown that pairs of synapses sharing the same presynaptic neuron and postsynaptic dendritic branch strongly correlate in size (***Figure 3***D) (***Motta et al., 2019; Bartol et al., 2015; Sievers et al., 2024***). We identified 37 synaptic pairs in L1 and 172 in L3 that share the same dendritic branch and the same presynaptic neuron (see Methods and Materials: Evaluating traces of plasticity on postsynaptic densities). To evaluate whether these pairs exhibit structural patterns consistent with correlation-based plasticity, we compared them to two control conditions: a shuffled dataset, in which synaptic pairs were randomly reassigned to eliminate any correlations, providing a lower bound (consisting of 1000 shuffled versions of the original pairs); and a sorted dataset, in which PSD areas were matched to their most similar counterpart to maximize pairwise size correlations, providing an upper bound (***Figure 3***E, Methods and Materials: Evaluating traces of plasticity on postsynaptic densities). The observed and shuffled data show low pair-size correlations (L1: *r* = 0.15, *p* = 0.35; L3: *r* = 0.18, *p* < 0.05; L1 shuffled: *r* = 0.00, *p* = 0.45; L3 shuffled: *r* = 0.00, *p* = 0.46). The sorted data shows strong pairsize correlations (L1 sorted: *r* = 0.98, *p* < 0.001; L3 sorted: *r* = 0.99, *p* < 0.001), consistent with prior reports in rat hippocampus (***Bartol et al., 2015***). Coefficient of variation (CV) analysis further supports this trend, only the sorted data differs significantly from the observed pairs, whereas the comparisons with the shuffled data do not reach significance (***Figure 3***F; L1 vs. L1 shuffled: *p* = 0.76; L1 vs. L1 sorted: *p* < 0.01; L3 vs. L3 shuffled: *p* = 0.14; L3 vs. L3 sorted: *p* < 0.001). Taken together, these results suggest that correlation-based plasticity does not leave strong anatomical signatures in this circuit and seems unlikely to be a primary driver of synaptic organization. However, its contribution cannot be ruled out through mechanisms not captured by static EM data, or obscured by concurrent structural plasticity (***Motta et al., 2019; Bartol et al., 2015; Ferreira Castro et al., 2023***).

### LN dendritic synaptic density is conserved throughout development

While our results suggest that synaptic scaling or correlation-based plasticity alone may not be the main drivers behind the observed synaptic sizes, this does not preclude other forms of synaptic regulation. One plausible candidate is synaptic density invariance, which we have shown in a companion paper to be observed across species (***Ferreira Castro and Cardona, 2024***), and may emerge through structural plasticity mechanisms (***Kirchner and Gjorgjieva, 2021; Kirchner et al., 2025; Lu et al., 2025***). We next investigated whether such density invariance is maintained in the developing circuit (***Figure 4***A).

**Figure 4.**
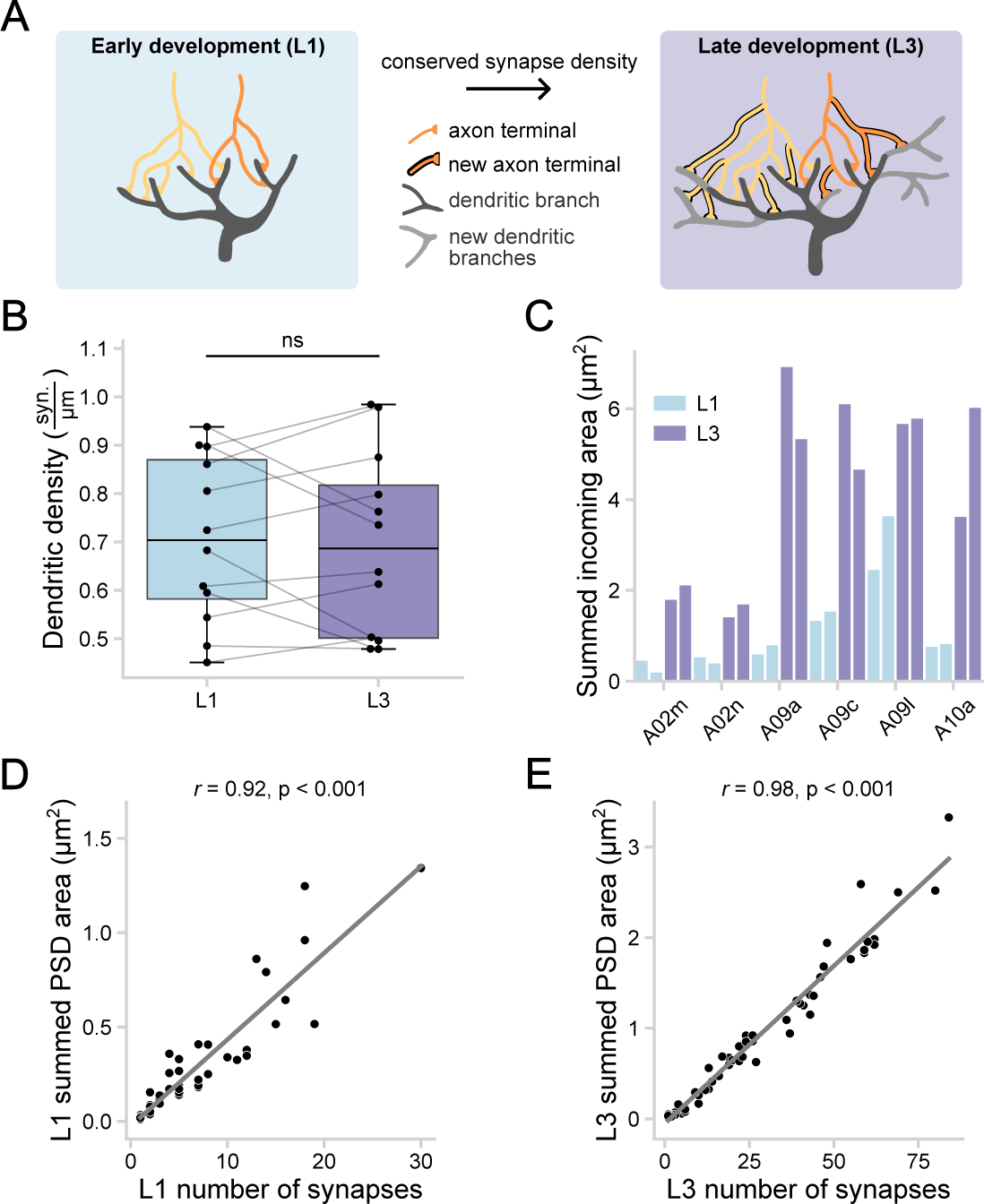
Maintenance of dendritic synaptic density throughout development. (**A**) Illustration of dendritic synaptic density maintenance during dendritic cable growth for the L1 (blue) and L3 (purple) developmental stage. Same color scheme is used in the remaining panels. (**B**) Dendritic synaptic density of LNs (one each in left and right side of the VNC) in L1 (*N* = 12) and L3 (*N* = 12). Lines connect individual LNs of L1 with their corresponding LN in L3. (**C**) Summed synaptic PSD areas of all the incoming mdIV synaptic connections to each LN in L1 and L3. Bars represent LNs of each specific type from the left and right sides of the VNC. (**D, E**) Correlation between the number of synapses (*N*_*L*1_ = 319; *N*_*L*3_ = 1562) between connected mdIV-LN neuron pairs and their summed synaptic PSD areas in L1 (*N*_*connection*_ = 48) (**D**) and L3 (*N*_*connection*_ = 64) (**E**).

To determine whether neurons regulate synapse number in proportion to dendritic growth, we quantified synaptic density across LN dendrites in both L1 and L3 and found no significant difference between the two developmental stages (***Figure 4***B; mean L1: 0.71 synapses per *μm*, mean L3: 0.70 synapses per *μm, p* = 0.86). Furthermore, individual LN synapse densities in L1 strongly correlate with their corresponding L3 counterparts (***Figure S2***A, *r* = 0.8, *p* < 0.01). These results suggest that neurons regulate synaptic numbers in proportion to dendritic growth, ensuring a stable synaptic density throughout development.

Theoretical models suggest that when synaptic density is maintained, uniform synapse sizes are sufficient to preserve neuronal excitability, reducing the need for precise size modulation (***Cuntz et al., 2021***). To test this, we asked how connection strength, defined here as the summed PSD area per mdIV-to-LN connection, changes across development. We observed a fivefold increase in summed PSD area of connected neurons between L1 and L3 (***Figure 4***C; ***Figure S2***B; dashed line, median fold change = 4.94). However, summed PSD areas increase linearly with synapse counts for each mdIV-to-LN connection (***Figure 4***D, E), with similar slopes in L1 (*r* = 0.92, *p* < 0.001, slope = 0.046) and L3 (*r* = 0.98, *p* < 0.001, slope = 0.035). These strong correlations suggest that connection strength is primarily determined by synaptic numbers, with average synaptic sizes per connection remaining relatively uniform (***Barnes et al., 2022***)(***Figure S2***B; solid line), supporting a density-conserving growth model.

Together, these findings support a developmental strategy in which circuit function is maintained by adding synapses in proportion to dendritic expansion rather than tuning synapse size. This mode of growth could ensure constant synaptic density, but raises the question of how such precise synaptic recruitment could be coordinated during neuronal growth (***Kirchner et al., 2025***).

### A mismatch of axonal and dendritic arbor growth across development

To understand how synaptic density is maintained during development, we next asked whether changes in the spatial arrangement of axons and dendrites, specifically, their physical overlap, play a role. Prior studies have shown that synaptic connectivity is strongly influenced by the proximity and geometry of axonal and dendritic arbors, and that even unspecific axonal and dendritic branching can lead to the emergence of appropriate connections in regions of neurite overlap (***Stepanyants and Chklovskii, 2005; Udvary et al., 2022; Moreno-Sanchez et al., 2024; Bird et al., 2021a***).

Building on this, we statistically verified that dendritic length and number of synaptic inputs scale proportionally across individual LNs (***Figure 5***A,B; *p* = 0.69; mean fold change cable length = 5.29, synaptic input = 5.18; ***Figure S3***A, *r* = 0.74, *p* < 0.01). This proportionality is expected assuming a growth program which preserves density. Despite this, mdIV axons exhibit a mismatch between structural growth and number of synaptic outputs: while presynaptic numbers in L3 is five times that in L1 (***Figure 5***C, D; mean fold change = 5.03), axon terminals are only 3.7 times longer in L3 compared to L1—a significantly smaller expansion (*p* < 0.05). We also found that the change in axon terminal length does not correlate with the increase in synaptic numbers (***Figure S3***B; *r* = 0.35, *p* = 0.49). These results indicate that presynaptic axons do not grow to match the scale of their postsynaptic targets, suggesting that the spatial proximity of axons and dendrites is not sufficient to explain how synaptic density is maintained during development.

**Figure 5.**
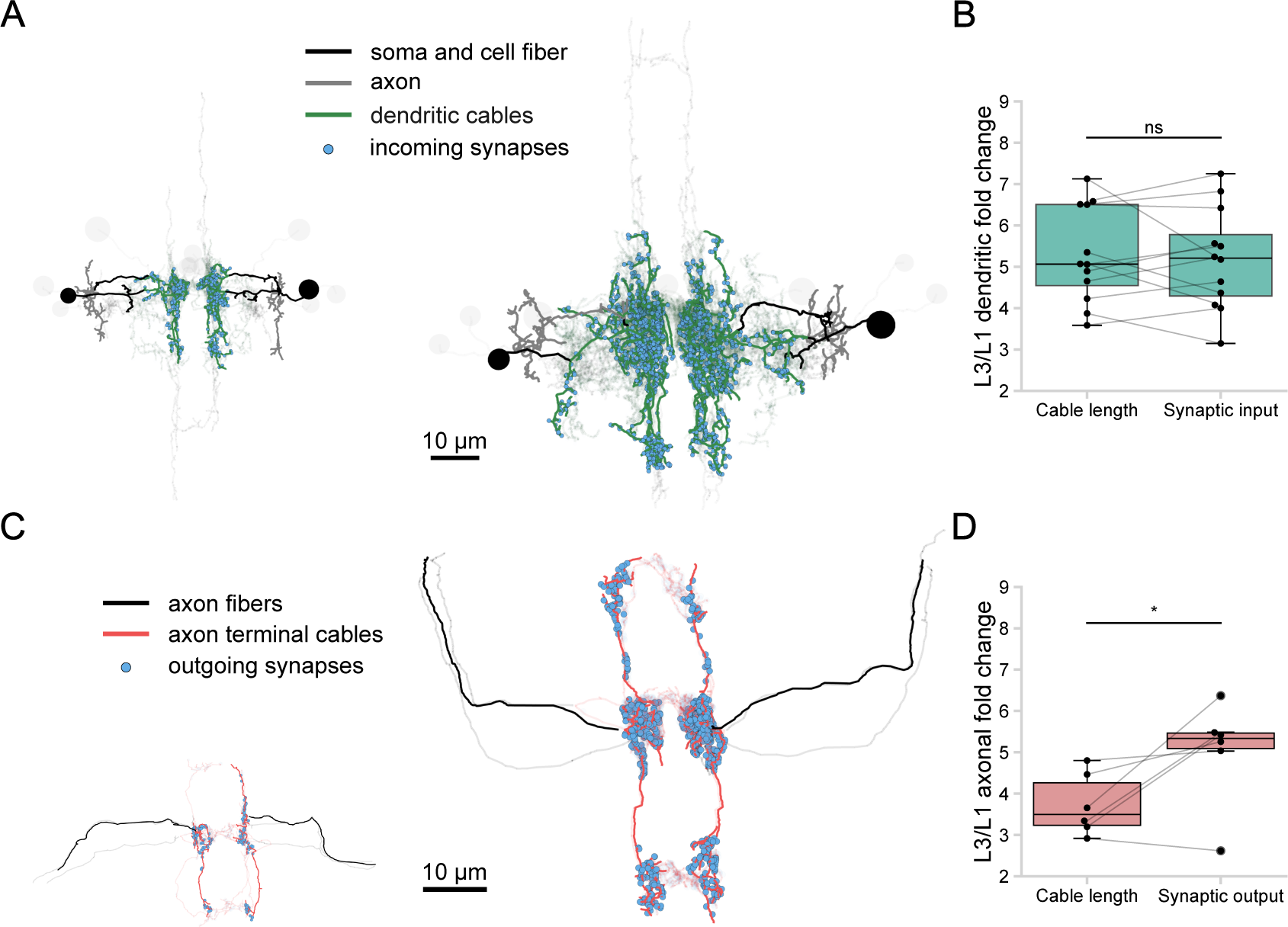
LN dendrite and mdIV axon scaling during postembryonic development. (**A**) Visualization of all LNs in L1 (left, *N* = 12) and L3 (right, *N* = 12), with two example LNs highlighted (A09l from the left and right side of the VNC), showing dendrites (green), axons (gray), and somas plus cell body fibers (black). Incoming synaptic connections to the dendritic trees are marked with blue dots. The remaining dendritic structures are shown with reduced opacity. An approximately fivefold increase in dendritic cable length and synaptic input is observed. (**B**) Fold change between L1 and L3 in dendritic cable length and dendritic synaptic input for all LNs (*N* = 12). Lines connect the cable and input ratios for each LN type. (**C**) Visualization of all mdIV axons in L1 (left, *N* = 6) and L3 (right, *N* = 6), with two example mdIVs highlighted (v’ada from left and right VNC), showing axon terminals (red) and remaining axonal cable (black). Outgoing synaptic connections from the axon terminals are marked with blue dots. All other axon terminals are displayed with reduced opacity. Visualization illustrates an approximately three-fold increase in axon terminal cable length and five-fold increase in axonal synaptic output. (**D**) Fold change between L1 and L3 in axon terminal cable length and axonal synaptic output for all mdIVs. Lines connect the cable and output ratios for each mdIV type.

### Increased presynaptic density compensates for reduced axon-dendrite overlap to conserve postsynaptic connectivity

Given the mismatch between dendritic and axonal growth, we asked how LN synaptic density might be conserved when spatial access to dendrites becomes limited. One possibility is that reduced axonal cable restricts synaptic opportunities, decreasing overall cable overlap. Alternatively, dendrites may compensate by growing more intermingled with axons, preserving synaptic density and connectivity despite smaller axonal expansion.

To test this, we developed a measure to quantify axon-dendritic cable overlap between neurons (Methods and Materials: Axodendritic cable overlap analysis). This measure identifies regions of dendritic cable that lie within a data-driven synaptic distance from axons (500 *nm*; Methods and Materials: Axodendritic cable overlap analysis), representing compartments where synapses could potentially form (***Figure 6***A). Quantifying axon-dendritic cable overlap reveals a significant ∼ 3-fold increase in overlapping dendritic regions (***Figure 6***B; mean mdIV to LN overlap L1 ∼ 50*μm*; L3 ∼ 150*μm, p* < 0.001), similar to axonal cable increase in development. However, when normalized by dendritic cable length, the proportion of dendrite accessible to axons reduces by ∼ 40% during development (***Figure 6***C, D; ***Figure S4***A; mean mdIV to LN overlap percentage L1 = 28%; L3 = 17%). This indicates that dendritic expansion outpaces axonal growth, reducing spatial access for synapse formation, potentially limiting connectivity preservation in the larval nociceptive circuit.

**Figure 6.**
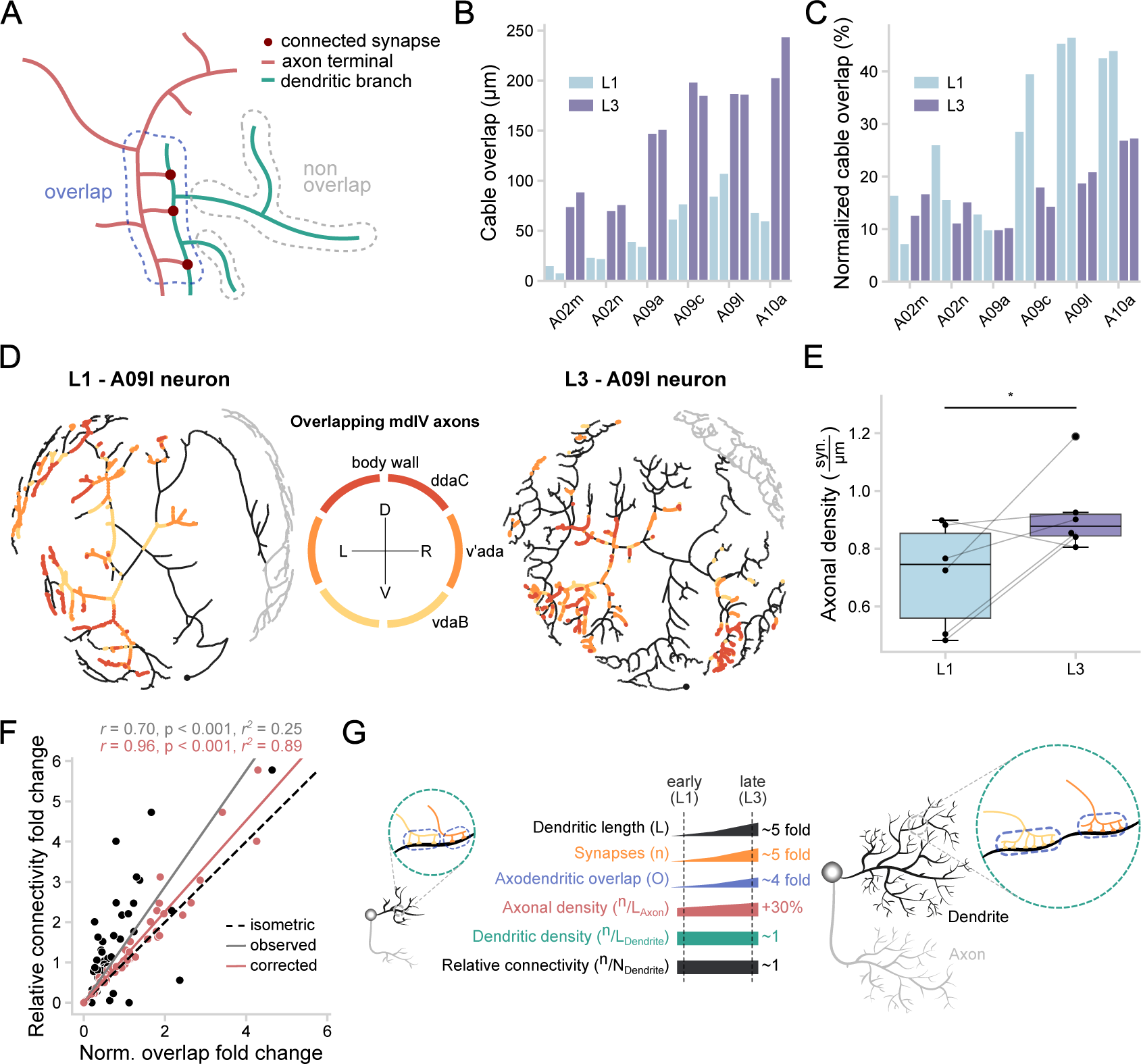
Presynaptic density compensates for reduced axon-dendrite overlap to conserve connectivity during development. (**A**) Schematic illustration of axon-dendritic overlap quantification between an mdIV axon and an LN dendrite. (**B**) Total axon-dendrite overlap length in L1 and L3 for all mdIV-to-LN connections. Bars represent LNs of each specific LN from the left and right sides of the VNC. (**C**) Normalized overlap, expressed as the percentage of dendritic cable in proximity to mdIV axons for each LN (100% = complete overlap). Bars same as in **B**. (**D**) Flattened topological visualizations of a representative LN (A09l, right hemisegment) in L1 and L3 using the scalable force-directed placement (SFDP) algorithm (***Hu***, ***2005***). Overlapping mdIV axons are color-coded by identity, illustrating the reduced normalized overlap between mdIV axons and the A09l LN in L3 compared to L1. (**E**) Axonal synaptic density for individual mdIV axons in L1 and L3; lines link matched axons across developmental stages. (**F**) Relationship between fold-change in axon-dendrite overlap and fold-change in connectivity for connections present at L1 and L3 (*N* = 48). The dashed line indicates a 1:1 isometric relationship expected if spatial overlap alone determines connectivity. The gray line shows the observed regression. Red points and fit show corrected predictions after adjusting overlap by the fold-increase in presynaptic density. (**G**) Schematic summary of identified circuit scaling properties in L1 (left) and L3 (right).

We next asked whether presynaptic terminals compensate for this reduction by increasing synaptic density within overlapping regions. To test this, we quantified axonal presynaptic density and found a significant ∼ 30% increase across development (***Figure 6***E; L1 mean = 0.71 synapses per *μm*; L3 mean = 0.92 synapses per *μm, p* < 0.05; mean fold increase = 1.37). In L1, axonal and dendritic synaptic densities were well matched, suggesting coordinated regulation of pre- and postsynaptic structure (*p* = 0.98; data from ***Figure 4***B and ***Figure S4***B). In L3, however, axonal density exceeds dendritic density (***Figure S4***B; *p* < 0.05), raising the question of whether the observed increase in axonal synaptic density is proportional to the reduction in axo–dendritic normalized overlap. To evaluate this, we scaled the L3 axonal density by the fold change in normalized overlap between L1 and L3 (Δ normalized overlap = 0.632). After this adjustment, we found that axonal and dendritic densities in L3 converge (***Figure S4***B; *p* = 0.18), suggesting that the apparent excess in axonal density reflects a compensatory increase in response to the reduced access of postsynaptic dendrite. We next asked whether the observed increase in presynaptic density also contributes to preserving relative connectivity at the level of individual mdIV-to-LN pairs. First, we calculated the fold-change in normalized overlap between L1 and L3 for each mdIV-to-LN pair and compared it to the corresponding fold-change in relative connectivity (***Figure 6***F). If spatial proximity alone dictates connectivity, these values would fall along the unity line (dashed line), indicating a one-to-one relationship. While normalized overlap and relative connectivity show significant correlation (*r* = 0.70, *p* < 0.001), relative connectivity is larger than expected based on overlap alone (*r*^2^ = 0.25, *slope* = 1.44; gray line, ***Figure 6***F). To account for this discrepancy, we adjusted the normalized overlap by the observed fold-increase in axonal synaptic density within the overlapping region for each connection (see Methods and Materials: Fold-change analysis of normalized axon-dendritic overlap and relative connectivity for analytical calculations). This adjustment yields a close match to the observed changes in connectivity (***Figure 6***F, red dots and line, *r* = 0.96, *p* < 0.001, *r*^2^ = 0.89, *slope* = 1.13), suggesting that modulating presynaptic density supports mdIV-to-LN relative connec-tivity conservation despite non-proportional growth of axons and dendrites.

Together, these findings support a growth strategy in which axons and dendrites may grow at different rates, yet compensatory mechanisms, specifically, increased presynaptic density, conserve stable connectivity during development (***Figure 6***G). While dendritic length scales approximately five-fold with larval growth from L1 to L3 (***Gerhard et al., 2017***), axon-dendritic overlap scales sublinearly, requiring a compensatory increase in synapse density along mdIV axons to maintain constant postsynaptic density. This adjustment enables synaptic input to track dendritic growth despite reduced overlap, ensuring stable relative connectivity across developmental stages.

### Maintenance of synaptic density and relative connectivity support functional stability across development

How do stable synaptic density and relative connectivity impact postsynaptic signal integration during development? To explore these relationships, we built passive single-cell biophysical models based on EM-derived morphologies and synaptic connectivity of LNs in L1 and L3 (Methods and Materials: Biophysical steady-state models of local neurons from the nociceptive circuit; ***Figure 7***A). This modeling approach is well suited for the electrotonically compact *Drosophila* neurons, as established by previous work in larval and adult nervous system (***Günay et al., 2015; Cuntz et al., 2013; Groschner et al., 2022***). Biophysical parameters were constrained from prior experimental measurements in *Drosophila* VNC neurons (***Günay et al., 2015; Cuntz et al., 2021***), and excitatory synaptic mdIV inputs strength matched the measured PSD distributions in this study (Methods and Materials: Biophysical steady-state models of local neurons from the nociceptive circuit; ***Figure 2***B).

**Figure 7.**
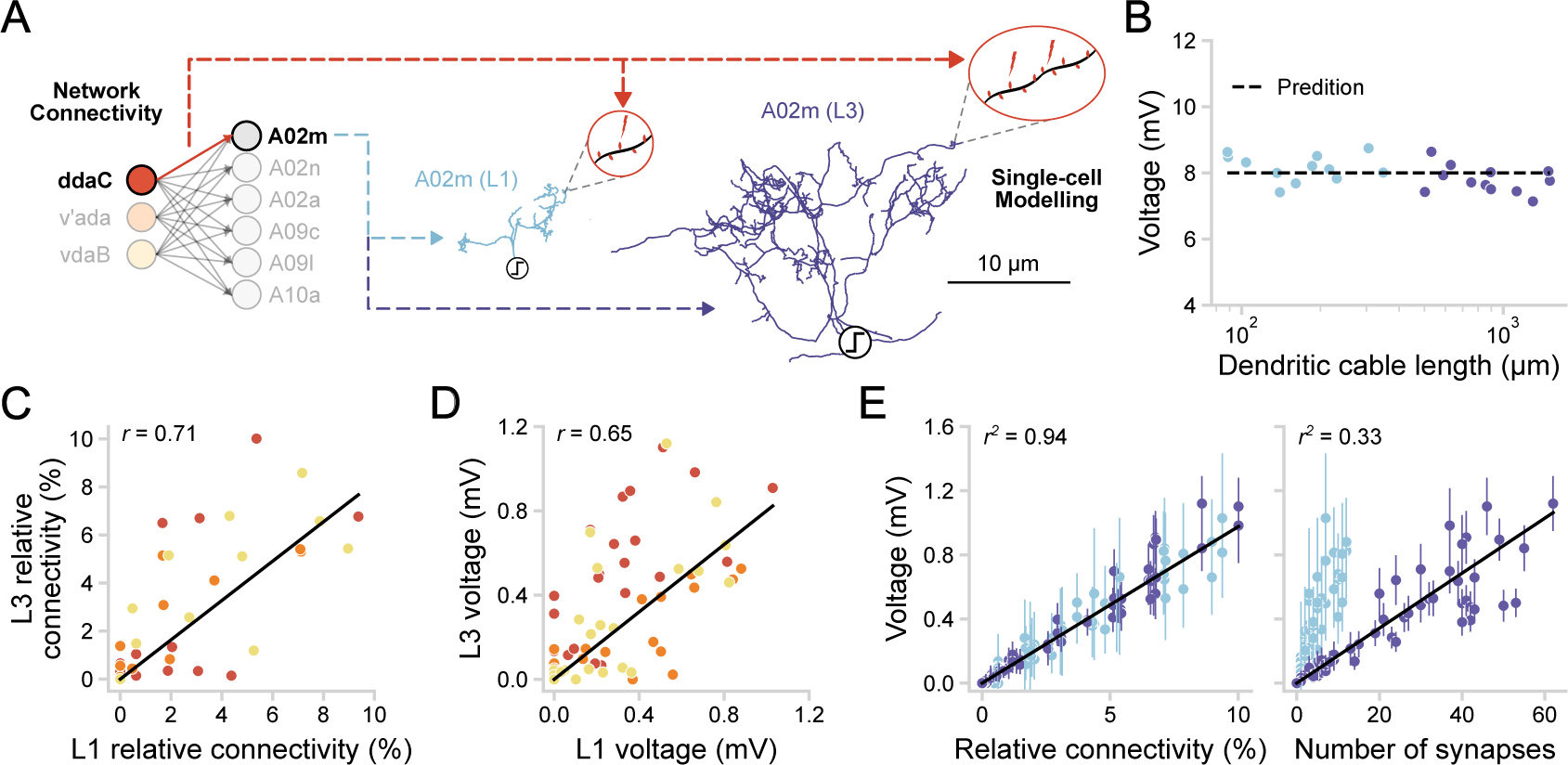
Synaptic density and relative connectivity conservation stabilize voltage responses across development. (**A**) Schematic representation of a pair of simulated LNs (A02m) across developmental stages. Inset highlights synapse activation with the same density and relative connectivity. Colors indicate developmental stages and presynaptic mdIV neuron identity, consistent with previous figures and applied throughout this figure. (**B**) Steady-state voltage responses to distributed synaptic inputs in all LN morphologies from the nociceptive circuit (*N*_total_ = 24, *L*1 = 12, *L*3 = 12), averaged over 1000 trials. Target voltage was set to *V*_*syn*_ = 8*mV*, to match spike threshold of VNC neurons from ***Günay et al. (2015***). The dashed line represents the analytical prediction, fitted using the same electrophysiological parameters as the models. (**C**) Relative connectivity of all mdIV-to-LN connections present in L1 and L3. The black straight line represents a linear regression fit (*r* = 0.94, *p* < 0.001). (**D**) Simulated voltage responses from all mdIV-to-LN connections across all neurons, averaged over 1000 trials. The black straight line represents a linear regression fit (*r* = 0.33, *p* < 0.001). Color scheme distinguishes presynaptic mdIV neurons as before. (**E**) Left: Simulated voltage responses from all mdIV-to-LN connections plotted against their corresponding relative connectivity as in (**C**). Error bars represent standard deviations. The black straight line represents a linear regression fit (*r*^2^ = 0.71, *p* < 0.001). Right: Similar plot, but with voltage responses plotted against the absolute number of synapses for the corresponding relative connectivity (*r*^2^ = 0.38, *p* < 0.001). Correlation coefficients (*r*) were calculated using Pearson correlation; *p*-values were obtained from permutation tests. Linear regression lines and coefficients of determination (*r*^2^) were included where applicable.

We first tested a theoretical prediction from the dendritic constancy principle. It states that steady-state voltage responses remain largely invariant to neuron size when synaptic density is constant and synaptic inputs are of similar strength (***Cuntz et al., 2021***). Specifically, the predicted voltage at the integration site (*V*_syn_) can be approximated as 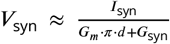, where *I*_syn_ is the total synaptic input current representing the sum of currents from distributed synaptic inputs, *G*_*m*_ is the membrane conductance reflecting the electrical properties of the dendritic membrane, *d* is the average dendritic tree diameter, and *G*_syn_ is the total synaptic conductance. To test this relation against biologically realistic conditions, we simulated steady-state voltage responses at the dendritic root (an anatomical proxy for spike initiation zone) to the activation of all synapses in our EM-constrained LN models in L1 and L3, and compared them to the analytical prediction computed using the same biophysical parameters as in the models (Methods and Materials: Biophysical steady-state models of local neurons from the nociceptive circuit). Despite more than an order-of-magnitude variation in dendritic length across L1 and L3, simulated voltage responses remained remarkably stable and closely matched the analytical predictions (normalized root-meansquare error *nRMSE* = 5.47%, relative to the predicted voltage responses; (***Figure 7***B). This agreement indicates that even in the presence of heterogeneous synaptic strengths, synaptic density conservation can maintain stable voltage responses across development.

We next asked whether relative connectivity also contributes to response stability at the level of specific mdIV-to-LN pairs (***Cuntz et al., 2021; Ferreira Castro and Cardona, 2024; Tobin et al., 2017***). Building on prior work showing conserved relative connectivity at the cell-type level during development (***Gerhard et al., 2017***), we verified the same relationship at the level of individual mdIV-to-LN connections between L1 and L3 (*r* = 0.71, *p* < 0.001; ***Figure 7***C) (***Gerhard et al., 2017***). To test the functional consequences of this correlation, we simulated responses to synaptic input from individual mdIV axons using our EM-constrained LN models. For each mdIV-to-LN pair, we used relative connectivity to determine the number of activated synapses at both developmental stages (Methods and Materials: Biophysical steady-state models of local neurons from the nociceptive circuit). We found that voltage responses for the same mdIV-to-LN connections are highly correlated across development (*r* = 0.65, *p* < 0.001), suggesting that relative connectivity helps maintain consistent postsynaptic responses for individual connections across development (***Figure 7***D).

Finally, we examined whether relative connectivity is a strong predictor of postsynaptic voltage responses. Plotting voltage response amplitude against relative connectivity reveals a strong linear relationship in both L1 and L3 (*r*^2^ = 0.91), whereas absolute synapse number shows a much weaker relationship (*r*^2^ = 0.38; ***Figure 7***E). This trend holds consistently across the full range of relative connectivity values (***Figure S5***). Together, these results suggest that voltage responses primarily reflect the proportion of active synaptic inputs, rather than the absolute number of synapses, and remain robust to changes in dendritic morphology and size during development (***Cuntz et al., 2021; Tobin et al., 2017***).

## Discussion

Our work reveals a strategy by which neurons can preserve functional stability across development. We generated the contactome of a fully mapped nociceptive circuit in the *Drosophila* larva using synaptic-resolution connectomic data, and quantified changes in synaptic size and connectivity between sensory mdIV neurons and downstream local neurons across two developmental stages: first and third instar. Despite growth, synaptic sizes remained largely stable, inconsistent with predictions of developmental synaptic upscaling. Additionally, we did not observe convergence in size for synapses sharing the same pre- and postsynaptic partners, suggesting that correlation-based plasticity does not shape synaptic sizes across the two developmental stages. We note that these inferences are based on the absence of anatomical signatures, and such processes may well be present but masked by other factors or remain undetectable with our current methods. Nonetheless, this suggests that these mechanisms are less likely to play a dominant role in maintaining circuit stability during postembryonic growth. Instead, we observed a striking invariance in postsynaptic density, indicating that LN dendritic growth is accompanied by proportional synaptogenesis to maintain density. However, this dendritic expansion is not matched by axonal growth, resulting in reduced axon–dendrite overlap during development—a mismatch compensated by increased presynaptic density. Computational modeling showed that conserved synaptic density invariance and relative connectivity, support stable postsynaptic voltage responses despite changes in size and morphology. Together, these findings provide a framework for how developing circuits preserve function amid substantial structural changes.

### Inferring developmental synaptic dynamics from static EM reconstructions

Our study provides a synaptic-level resolution analysis of the *Drosophila* nociceptive circuit across development, but EM offers only a static snapshot of synaptic architecture (***Motta et al., 2019; Bartol et al., 2015; Ferreira Castro et al., 2023; Barnes et al., 2022***). Dynamic processes such as short-term plasticity, neuromodulation, and intrinsic excitability, which modulate synaptic integration, are not captured in our dataset but likely complement the developmental structural stability we describe (***Gorur-Shandilya et al., 2020; Bargmann, 2012; Ripoll-Sánchez et al., 2023***). Additionally, because EM imaging requires separate specimens, our L1 and L3 datasets come from different individuals rather than being a longitudinal progression within the same animal(***Gour et al., 2021; Yim et al., 2024; Witvliet et al., 2021***). Future light microscopy studies using longitudinal time-lapse imaging could complement our work by tracking structural remodeling within the same organism, albeit at lower resolution (***Ferreira Castro et al., 2020; Shree et al., 2022; Baltruschat et al., 2020; Tripodi et al., 2008; Heckman and Doe, 2022; Rigaux et al., 2025***). Despite these limitations, the key feature we observe, invariant synaptic density across development, is also found across other species, as shown in our companion study (***Ferreira Castro and Cardona, 2024***). This cross-species consistency indicates that our findings reflect conserved biological features rather than artifacts of a particular dataset. Moreover, our computational modeling suggests a potential functional role for this density invariance in maintaining stable postsynaptic responses across development.

An assumption in our study is that PSD area serves as a proxy for synaptic strength. In mammals, PSD size strongly correlates with synaptic conductance and functional strength (***Cheetham et al., 2014; Holderith et al., 2012; Holler et al., 2021***), however, this relationship is more complex in *Drosophila*. Presynaptic properties, such as, vesicle release probability and active zone size, can also be major determinants of synaptic efficacy (***Davis and Bezprozvanny, 2001; Paradis et al., 2001; Murthy et al., 2001; Turrigiano, 2012***). This raises the possibility that PSD size alone may not capture synaptic strength across species equally well (but see ***Davis and Goodman (1998); DiAntonio et al. (1999***)). However, in insects, the prevalence of polyadic synapses—where multiple postsynaptic neurons share the same presynaptic site—imposes a structural constraint that limits presynaptic release (***Meinertzhagen and O’neil, 1991***). This suggests that, despite its limitations, PSD still is the best structural correlate of connection strength in *Drosophila* (***Barnes et al., 2022***). Combining electrophysiological recordings with EM reconstructions in future correlative studies will be necessary to establish a clearer link between PSD size and synaptic efficacy in *Drosophila*. Importantly, our computational modeling demonstrates that voltage response stability is robust to variability in synaptic strengths. Therefore, even if PSD size were an imperfect proxy for connection strength, our core conclusion that synaptic density invariance and relative connectivity preserve circuit function during growth, would still hold (***Figure 7***).

### Assumptions in EM-constrained modeling

Our computational models, while providing valuable insights into how synaptic density invariance contributes to functional stability, necessarily rely on simplifying assumptions A key assumption is that all neurons have passive, homogeneous electrophysiological properties. This is biologically justified in modeling *Drosophila* neurons, as they often behave passively due to their small size and electrotonic compactness (***Abdelrahman et al., 2021; Cuntz et al., 2013; Groschner et al., 2022; Cuntz et al., 2021; Günay et al., 2015***). In our models, dendritic membrane conductance was constrained with experimental measurements of motor neurons in the larval VNC because comparable data for LNs are not available (***Günay et al., 2015***). This choice is reasonable because the passive membrane properties of small, electrotonically compact cells are broadly conserved across neuron types in *Drosophila* (***Günay et al., 2015; Gouwens and Wilson, 2009***).

Additionally, our model only includes excitatory synapses, due to the fact that allmdIV-LN connections within the studied circuit are excitatory. As a result, they do not account for the potential modulatory effects of inhibitory inputs that may influence circuit function at a broader network level(***Ohyama et al., 2015; Jovanic et al., 2016***). We also assumed uniform synaptic kinetics, conductances, and reversal potentials and only calculated the steady-state response, which simplifies synaptic diversity and integration properties in the biological system (***Cuntz et al., 2021***). Future work incorporating heterogeneous synaptic properties and time-resolved synaptic integration will further our understanding of how connectivity and excitability are maintained across developmental scales.

### Synaptic density invariance and circuit stability-plasticity dilemma across development

Our findings suggest that conserving synaptic density is key to maintain circuit stability during development. Activity-dependent changes in synaptic size appear to play a lesser role, consistent with prior computational models of invertebrate motor circuits (***Gjorgjieva et al., 2016***). In contrast, in mammals, activity-dependent synaptic plasticity mechanisms, such as Hebbian plasticity, play a much bigger role in refining synaptic connections to achieve circuit assembly as shown both experimentally and in computational models (***Miller et al., 1989; Fumarola et al., 2022; Bartsch and van Hemmen, 2001; Kirkwood and Bear, 1994; Richter and Gjorgjieva, 2017***). Even so, homeostatic mechanisms are needed to regulate the positive feedback instability generated by Hebbian synaptic plasticity (***Van Rossum et al., 2000; Wu et al., 2020; Zenke et al., 2017***). Hence, while Hebbian mechanisms are important for synaptic size refinement, other mechanisms might be need to maintain stability once a network is established (***Fox and Stryker, 2017; Keck et al., 2017; Toyoizumi et al., 2014; Motta et al., 2019; Sievers et al., 2024***). For example, mammalian dendritic trees also employ structural homeostasis to conserve synaptic numbers and whole-cell input strength after Hebbian-like plasticity, suggesting a dynamic balance between synaptic plasticity and density conservation (***Royer and Paré, 2003; Bourne and Harris, 2011; Eckmann et al., 2024; Kirchner et al., 2025; Field et al., 2020***).

Our results also suggest that synaptic scaling alone is unlikely to account for functional stability during development in the larval nociceptive circuit. From a theoretical standpoint, synaptic up-scaling could offset the biophysical consequences of neuronal growth without requiring changes in synapse number (***Turrigiano, 2008***). Although we cannot rule out the possibility that upscaling effects are distributed in ways that escape detection with our current approach, it may also reflect structural or metabolic constraints. Simply increasing synaptic size while maintaining synapse number during body expansion could be detrimental to circuit design. First, synaptic growth incurs supralinear energy costs, making unrestricted expansion metabolically inefficient (***Sterling and Laughlin, 2015***). Second, geometric constraints impose spatial limitations, as excessively large synapses may disrupt optimal structural packing and increase conduction delays (***Stepanyants and Chklovskii, 2005; Chklovskii et al., 2002; Piazza et al., 2025***). Thus, rather than relying solely on global synaptic scaling, circuits may prioritize maintaining synaptic density and size within a regulated range to ensure functional stability, as also shown across species in our companion paper (***Ferreira Castro and Cardona, 2024***).

The conservation of synaptic density and relative connectivity across development raises the question of which underlying mechanisms control synaptic inputs during body growth. Synapse formation is thought to result from a combination of factors, including spatiotemporal partner availability, branching dynamics, and molecular recognition cues between pre- and postsynaptic cells (***Kiral et al., 2021; Agi et al., 2020, 2024; Valdes-Aleman et al., 2021; Hoersting and Schmucker, 2021; Ferreira Castro et al., 2020; Barabási et al., 2025***). This framework partially aligns with the developmental scaling of relative connectivity found in our study: as new stable axonal and dendritic branches are formed during development, synaptic recruitment from preferential partners increases accordingly (***Kiral et al., 2021***).However, these processes alone cannot fully account for the conservation of synaptic density and input fractions during growth. Our data and analytical calculations suggest that regulating axonal presynaptic density is necessary to maintain dendritic synaptic density and relative connectivity (***Elkahlah et al., 2020; Petersen et al., 1997***). Future studies should continue to investigate the functional implications of manipulating the spatiotemporal availability of presynaptic partners, and how this affects the relative connectivity of postsynaptic inputs (***Valdes-Aleman et al., 2021; Kiral et al., 2021***). Furthermore, glia-neuron interactions may contribute to maintaining synapse numbers, as they are uniquely positioned in close proximity to both pre- and postsynaptic sites (***Heckman and Doe, 2021***). In vertebrates, glial cells such as microglia and astrocytes regulate synaptic homeostasis through pruning and maintenance (***Wu et al., 2024; Perez-Catalan et al., 2021; Clarke and Barres, 2013***). Although *Drosophila* glia are less well understood, their interactions could similarly help preserve synaptic density and relative connectivity(***Heckman and Doe, 2021; Ackerman et al., 2021; Yildirim et al., 2019; Bittern et al., 2021***).

### Relative connectivity conservation and the maintenance of circuit function

The consistent regulation of relative connectivity across species and brain regions suggests it is an important feature of circuit organization (***Gour et al., 2021; Schneider-Mizell et al., 2025; Schlegel et al., 2024; Winding et al., 2023***). Rather than hardwiring specific synaptic contacts, neural circuits appear to be organized through tight control of relative connectivity between synaptic partners, which may help explain variability in absolute synaptic counts even among genetically identical individuals (***Witvliet et al., 2021; Schlegel et al., 2024***). Why might brains prioritize conserving relative connectivity over precise synaptic numbers across development? The answer is twofold. First, the dendritic constancy principle predicts that, for a fixed synaptic density, postsynaptic voltage responses scale with the fraction of active synapses relative to the total inputs (***Cuntz et al., 2021***). This property is independent of neuron size and connectivity, effectively normalizing connection strength as neurons grow (***Bird et al., 2021b***). Thus, preserving relative connectivity during development functions as a form of weight normalization that stabilizes neuronal output despite large changes in size.

Second, stochastic developmental processes (***Linneweber et al., 2020; Hiesinger and Hassan, 2018; Barabási et al., 2025***) and environmental factors contribute to variability in brain wiring, even as circuits maintain stable functional output (***Kiral et al., 2021; Gerhard et al., 2017***). Hardwiring exact synapse numbers would require extreme wiring precision and could lead to catastrophic failures in noisy, dynamic systems (***Hiesinger and Hassan, 2018***). In contrast, encoding relative connectivity allows neurons to adapt to diverse developmental conditions while preserving functional responses (***Tobin et al., 2017; Ferreira Castro and Cardona, 2024***). This strategy is particularly relevant for maintaining circuit stability amid developmental noise, including variation in partner availability (***Kiral et al., 2021***), and asymmetries in body growth rates, such as left-right brain growth disparities (***Pedigo et al., 2023***). Additionally, it may buffer against external perturbations, such as food source scarcity (***Mirth and Riddiford, 2007***) or temperature fluctuations (***Kiral et al., 2021***), which influence growth trajectories.

## Conclusion

By integrating synaptic-resolution EM data and single-cell computational modeling within a well-defined system, our results show that the conservation of synaptic density and relative connectivity can support stable neuronal responses during development, despite dramatic growth and expansion. As advances in synaptic-level resolution connectomics continue to expand the range of mapped nervous systems (***Dorkenwald et al., 2024; Schneider-Mizell et al., 2025; Sievers et al., 2024; Velicky et al., 2023; Petkova et al., 2025; Shapson-Coe et al., 2024***), integrating relative connectivity analyses may open new avenues for uncovering general principles of neural computation across species and developmental timescales.

## Acknowledgments

This work was supported by the Daimler and Benz Foundation (to AFC), a project in the NeuroMac consortium (CRC/TRR 167 to JG), and core funding from the Technical University of Munich (TUM). In ***Figure 1***A and B, icons downloaded from bioRender have been used (wasp and larva icons). We thank all members of the ‘Computation in Neural Circuits’ group and Elizabeth Herbert and Matt Getz for useful discussions and comments on the manuscript. We also thank Chris Barnes and Dylan Festa for their support in setting up BigCat and managing the connectomics datasets. We thank Albert Cardona for helpful discussions and his support during this project.

## Author contributions

Ingo Fritz, Data curation, Investigation, Visualization, Formal analysis, Methodology, Software, Writing – original draft, Writing - review and editing; Feiyu Wang, Data curation; Ricardo Chirif Molina, Data curation; Nikos Malakasis, Visualization, Supervision; Julijana Gjorgjieva, Conceptualization, Supervision, Funding acquisition, Investigation, Methodology, Project administration, Writing – review and editing; André Ferreira Castro, Conceptualization, Data curation, Investigation, Visualization, Formal analysis, Methodology, Software, Supervision, Project administration, Writing – original draft, Writing - review and editing, Funding acquisition.

## Data and code declaration

Availability of data and materials: all data and code are publicly available on GitHub: https://github.com/comp-neural-circuits/circuit-stability-development.

## Methods and Materials

### Toolboxes and Libraries

In this study, all analyses were conducted using Python (version **3.12.7**). The scripts used for data processing, analysis, and figure generation are publicly available on GitHub: https://github.com/comp-neural-circuits/circuit-stability-development. We obtained two connectomes (first and third instar larva) from CATMAID (Collaborative Annotation Toolkit for Massive Amounts of Image Data, https://catmaid.readthedocs.io/en/stable/), where the connectomics data is accessible through the Virtual Fly Brain platform. Specifically, we used the L1 connectome (https://l1em.catmaid.virtualflybrain.org/) and the L3 connectome (https://l3vnc.catmaid.virtualflybrain.org/) in this work. Simulations were performed using the TREES Toolbox (MATLAB, https://www.treestoolbox.org/).

To load and analyze the connectomes, we used PyMaid (a Python interface for CATMAID, https://pymaid.readthedocs.io/en/latest/), which facilitates interaction with CATMAID servers via Python scripts. For additional analysis and visualization of neurons, we utilized the NAVis package (Neuron Analysis and Visualization, https://navis-org.github.io/navis/). All EM-derived data—including skeletons, synapse annotations, and soma coordinates—were imported into analysis environments using NAVis (Python, https://github.com/navis-org/navis), natverse (R, (***Bates et al., 2020***)), and the TREES Toolbox (MATLAB, https://www.treestoolbox.org/).

### Electron microscopy data and morphologies

Neuronal morphologies were obtained from two electron microscopy volumes of the peripheral nervous system in *Drosophila melanogaster* larvae, representing two distinct developmental stages. The first dataset consists of 6 mdIV axons and 12 local neurons (LNs) from the A1 segment of a first instar (L1) larva (***Ohyama et al., 2015; Jovanic et al., 2016; Gerhard et al., 2017***). The second dataset contains 6 mdIV axons and 12 LNs from the A3 segment of a third instar (L3) larva (***Gerhard et al***.,***2017***). These datasets were selected because they provide dense, expert-annotated reconstructions of the larval nociceptive circuit, enabling direct comparative analysis across developmental stages.

EM skeletons for both L1 and L3 were downloaded from the Virtual Fly Brain platform (https://virtualflybrain.org) using custom Python and R scripts. All coordinates were isotropically rescaled to 1× 1 × 1*μm* resolution. Skeleton statistics and morphometric measurements were computed using custom Python, R, and MATLAB scripts within the respective environments. All analysis scripts and data files related to morphology and synapse locations will be made publicly available upon submission.

### Post synaptic density (PSD) segmentation and measurement

PSD area was used as a proxy for synaptic strength between mdIV axons and LNs. We adopted the volumetric annotation approach described in ***Barnes et al. (2022***), where PSDs were manually segmented in each EM slice containing a synaptic contact. Synaptic locations were extracted using CATMAID’s API (***Saalfeld et al., 2009***), and the surrounding image data were stored in extended CREMI-format HDF5 files. PSDs were identified as membrane regions adjacent to the synaptic cleft and exhibiting postsynaptic specializations. Using BigCat (https://github.com/saalfeldlab/bigcat), a volumetric annotation tool, PSDs were annotated slice by slice, skeletonized, smoothed, and converted into surface area estimates by multiplying 2D arc lengths with the z-resolution (50*nm*). The full 3D synaptic area was computed as the sum of these estimates across z-sections. For details of this procedure, see ***Barnes et al. (2022***).

In total, we measured PSD areas for 319 synapses in L1 and 1562 synapses in L3, capturing all identified mdIV-to-LN connections. To visualize PSD area distributions, we log-transformed the measurements. Distributional differences between L1, L3, and scaled L1 data were evaluated using two-sample Kolmogorov–Smirnov (KS) tests (***Figure 2***B; ***Figure 2***C).

### Evaluating traces of plasticity on postsynaptic densities

To test whether postsynaptic density (PSD) areas reflect patterns consistent with correlation-based plasticity, we conducted two complementary analyses: (1) rank-based analysis of connectivity strength, (2) PSD similarity among synaptic pairs sharing the same pre- and postsynaptic dendritic branch.

For each LN, we ranked its connected mdIV neurons based on the number of synapses they formed with that LN. The mdIV neuron with the highest number of synapses was assigned rank 1; the neuron with the fewest was assigned rank *N*, where *N* is the total number of connected mdIVs. Ties were resolved by random assignment. For each rank, we computed the mean number of synapses and the mean PSD area of the corresponding mdIV-to-LN connections. These were plotted as a function of rank (***Figure 3***B, C). To assess whether synaptic size scales with connectivity rank, we computed the Pearson correlation coefficient between mean PSD area and rank.

To examine whether individual synaptic pairs reflect correlation-based plasticity, we isolated cases in which exactly two synapses from the same mdIV axon contacted the same LN dendritic branch. Dendrites were separated at branch points, and pairs were selected only if exactly two synapses from a given mdIV neuron innervated a given branch. This yielded 37 valid synaptic pairs in L1 and 172 in L3, following criteria established in ***Bartol et al. (2015***).

We assessed PSD area similarity within each pair using two metrics: the Pearson correlation coefficient and the coefficient of variation (CV). To benchmark these values, we generated two control datasets: (1) 1000 shuffled control in which PSD areas were randomly reassigned to all pairs, and (2) a sorted control in which PSDs were artificially paired to maximize similarity (e.g., the two largest PSDs were paired, and so on). We computed the same statistics for each control ***Figure 3***E, F). CV comparisons between conditions were assessed using two-sample t-tests, corrected for multiple comparisons using Bonferroni adjustment (n=2; ***Figure 3***F).

### Separation of axonal and dendritic compartments

To distinguish axonal and dendritic compartments within each neuron, we used the navis.morpho.split_axon_dendrite function from the NAVis package. This function separates neuronal arborizations based on either bending-flow or synapse-flow centrality ***Schneider-Mizell et al. (2016***). These methods allow compartmentalization based on the geometric structure of the neuron or on the spatial distribution of synaptic inputs and outputs. The latter approach is particularly relevant for local neurons (LNs), which exhibit clear spatial segregation between pre- and postsynaptic sites, in contrast to mdIV axons, which are dominated by presynaptic outputs and contain relatively few postsynapses.

LN skeletons were split using synapse-flow centrality, leveraging the positions of annotated pre- and postsynapses. We also separated axons and dendrites from the primary cell body fiber (including the soma), which lacks synapses and simply links the soma to the neuropil arborizations. This distinction enabled dendrite-specific morphometric analyses, including the computation of synaptic density (***Figure 4***B), dendritic cable length (***Figure 5***B, D), and axodendritic overlap with mdIV inputs (***Figure 6***B, C, D, F).

For mdIV axons, we used the bending-flow method to segment axonal fibers from axon terminals based on geometric inflection points and branch trajectories. This procedure reliably identified the point at which each axon bends upon entering the neuropil and begins branching to form synaptic terminals. We excluded the axonal fiber from calculations of terminal cable length (***Figure 5***C, D) and synaptic density (***Figure 6***E), since this compartment lacks synapses and does not contribute to output strength or potential connectivity (***Figure 5***C) (***Schneider-Mizell et al., 2016***).

### Relative connectivity calculation

To quantify the relative contribution of each mdIV neuron *i* to the total synaptic input of a given LN *j*, we computed the fraction of synaptic input that neuron *i* contributes. Specifically, the number of synapses in each connection *n*_*ij*_ was divided by the total number of incoming synapses of presynaptic partners *k* to LN *j*, denoted *N*_dend,*j*_ =Σ_*k*_*n*_*kj*_:

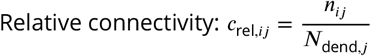

This measure reflects the proportion of synaptic input contributed by mdIV neuron *i* to LN *j*.

Relative connectivity has been shown to be largely conserved between L1 and L3 datasets (***Gerhard et al., 2017***), and thus serves as a proxy for functionally stable input structure during development.

We used this metric to evaluate whether developmental changes in axodendritic geometry and synapse number preserve relative input proportions. Specifically, we compared relative connectivity values to axon-dendritic overlap scaling measurements, testing whether isometric expansion alone could account for the observed connectivity patterns, or whether additional mechanisms, such as, axonal synapse density increases, contribute to preserving relative connectivity (***Figure 6***F, ***Figure 7***C, ***Figure S5***A, B).

### Axodendritic cable overlap analysis

To quantify potential synaptic conctacts between mdIV axons and LN dendrites, we measured axodendritic cable overlap based on spatial proximity between skeletonized arbors. All neurons were first resampled to a uniform resolution of 0.1 *μm* (i.e., 10 nodes per *μm*) to ensure consistent node density for overlap computations.

For each mdIV–LN pair, we evaluated which dendritic nodes lay within a 500 *nm* Euclidean radius of any mdIV axon node. This 500 *nm* threshold was chosen based on empirical measurements of the Euclidean distance between presynaptic and postsynaptic skeleton nodes in our dataset: we computed the median of these distances and added two interquartile ranges to define a conser-vative, biologically plausible proximity bound (***Bates et al., 2020***). This value aligns with distances typically observed in *Drosophila*.

Distances were computed using Euclidean distance between resampled skeleton nodes (not mesh models or voxel volumes). Each dendritic node was labeled as overlapping if it lay within 500 nm of any axonal node. This overlap metric is directional: we assessed whether dendritic cable was within 500 nm of axons, but not the reverse. All dendritic nodes within this threshold were considered, not just the closest. Overlap was defined as the total dendritic cable length of a given LN *j* that lies within a biologically plausible synaptic distance (500 nm) from a presynaptic mdIV axon *i*, yielding the absolute overlap (***Figure 6; Figure S4***):

*O*_*ij*_ = length of dendrite *j* within 500 nm of axon *i*

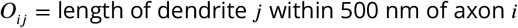

To account for differences in dendritic arbor size, we normalized overlap by the total dendritic length:

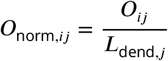

This represents the proportion of dendrite accessible to a given axon and serves as a proxy for spatial opportunity to form synapses. To avoid double-counting in aggregate analyses, we ensured multiple axons overlapping the same dendritic segments were only counted once in total overlap estimates (***Figure 6***B, C).

### Fold-change analysis of normalized axon-dendritic overlap and relative connectivity

To test whether spatial proximity alone predicts synaptic connectivity across development, we computed the normalized axon-dendritic cable overlap *O*_norm,*ij*_ between each mdIV-to-LN pairs. We further refined this quantity by incorporating local presynaptic density information. Let *n*_*ij*_ be the number of synapses made by axon *i* onto dendrite *j*. The local axonal synaptic density in the overlapping cable is:

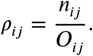

Multiplying normalized overlap by this presynaptic density gives the corrected normalized overlap:

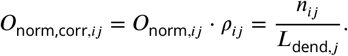

This quantity expresses the number of synapses from axon *i* onto dendrite *j* per unit length of the postsynaptic arbor. To assess whether this corrected overlap reflects synaptic contacts, we compared it to the relative connectivity:

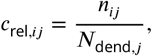

where *N*_dend,*j*_ is the total number of synapses received by dendrite *j* from all presynaptic partners. For the relationship *O*_norm,corr,*ij*_ ≈ *c*_rel,*ij*_ to hold across development, that is, an isometric relationship, dendritic synaptic density must remain approximately constant:

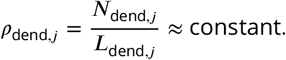

We empirically validate this assumption (***Figure 4***). From this, we rearrange:

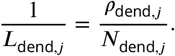

We can then substitute:

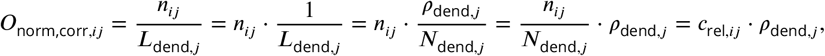

showing that corrected normalized overlap is proportional to relative connectivity, scaled by dendritic synaptic density. Because *ρ*_dend,*j*_ is constant across development, the scaling factor cancels out in fold-change (Δ) comparisons:

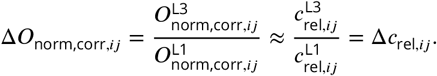

Empirically, we observed that:

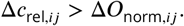

This indicates that normalized overlap alone underestimates the increase in relative connectivity between L1 and L3. Thus, specifically, from the equation

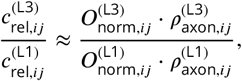

we can infer that the missing factor is the increase in presynaptic density:

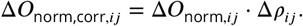

This shows that relative connectivity can only scale more steeply than normalized overlap if axonal synaptic density increases over development. Moreover, because postsynaptic synapse density remains approximately constant across stages 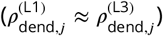, changes in *ρ*_axon,*ij*_must fully accountfor the discrepancy between overlap-based and observed connectivity changes. Thus, the correction based on presynaptic density is not only justified but necessary to reconcile developmental shifts in morphology with preserved connectivity structure.

In our data, applying this correction brought predicted and observed connectivity into close alignment (*r* = 0.96, *r*^2^ = 0.89), demonstrating that presynaptic density modulation compensates for reduced overlap and is sufficient to preserve relative connectivity during development (***Figure 6***F, G). This analysis underscores the importance of stable dendritic synaptic density across development. Without this conservation, the correction would not recover relative connectivity, and spatial overlap alone would fail to explain observed connectivity patterns.

### Biophysical steady-state models of local neurons from the nociceptive circuit

To investigate the role of synaptic density and relative connectivity conservation in maintaining functional responses across developmental stages, we developed passive steady-state models for all LN neurons in our dataset. We used .swc files derived from EM-based 3D reconstructions of LN dendrites from first (L1) and third (L3) instar stages. All morphologies were imported into the TREES Toolbox (MATLAB) using the load_tree function and resampled at 10*nm* resolution (***Cuntz et al., 2010***). Cable properties were defined with a constant dendritic diameter of *d* = 0.78 *μ*m and membrane conductance *G*_*m*_ = 38 *μ*S/cm^2^, consistent with larval ventral nerve cord (VNC) neurons (***Günay et al., 2015***).

Synapses were placed uniformly across the dendritic arbor in each trial. The number of synaptic counts per neuron was determined from the mean EM-derived synapse density at L1 and L3 in each trial (*ρ* ≈ 0.7). The total synaptic input current (*I*_syn_) was estimated using the dendritic con-stancy equation, *V*_syn_ = *I*_syn_/(*G*_*m*_ · π · *d* + *G*_syn_), with *G*_syn_ initially omitted to compute the total input required to reach a target steady-state voltage of approximately 8 mV (***Günay et al., 2015***). *G*_syn_ was initially omitted because it is itself a function of the total synaptic current, rendering the full expression implicit; solving for *I*_syn_ therefore requires an initial estimate under passive (conductance-free) conditions. The 8 mV target was chosen to reflect the approximate spiking threshold of local interneurons in the larval VNC (***Günay et al., 2015***). To compensate for the reduced input resistance caused by synaptic conductances, we introduced a small voltage overshoot (Δ*V*) to the target voltage during initial current estimation. This overshoot was computed using the compute_overshoot function, which numerically solves for the minimal Δ*V* required for a conductance-based synaptic input to reach the target voltage (8 mV), based on membrane parameters and synaptic density. Total synaptic conductance was computed as 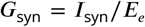, using a reversal potential of *E*_*e*_ = 40 mV (***Cuntz et al., 2021***). This conductance was divided among all synapses, with individual conductances being drawn from a log-normal distribution and scaled such that their sum matched the calculated 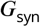 mimicking measured synaptic sizes. All simulations were then performed using the syn_tree from the TREES Toolbox, as described previously (***Cuntz et al., 2021***).

In our models, to assess how voltage responses scale with the proportion of active inputs, we varied the percentage of activated synapses according to empirically measured relative connectivity values for each mdIV-to-LN pair. To reduce variability arising from sampling differences between individual left and right hemisegments in our connectomes, we averaged each mdIV-to-LN connection across its bilateral symmetric counterparts and used this average as a single representative measure of relative connectivity (***Figure 7***C). This symmetrized metric was then used for all subsequent comparisons in the single-cell models, ensuring a more robust estimate of developmental connectivity patterns. For each simulation, the neuron’s voltage response to activated synapses was simulated 1000 times, and the results were averaged to obtain stable estimates. To increase biological realism, synaptic densities were varied around the mean using the empirically observed standard deviation across measured densities. The number of synapses activated in each trial was derived from relative connectivity, defined as 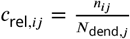, where larger dendritic arbors—despite having similar synaptic densities (centered around the observed mean of ∼ 0.7 synapses/*μ*m)— receive more synapses overall due to their size. In all simulations, activated synaptic locations were independently resampled at each trial to account for spatial location variability. Voltage responses were calculated at the root node using the syn_tree. Final outputs using the syn_tree function were the steady-state membrane potentials (*V*_syn_) at the dendritic root, representing the integrated synaptic input under passive conditions.

Finally, we also tested extreme activation patterns by selectively stimulating either the most distal or the most proximal synapses (***Figure S5***). Activation levels were incrementally increased in even steps from 0% to 100% of synapses for each condition, allowing us to assess how input location and coverage affect voltage responses across the full activation range. Functional responses from these simulations were averaged across all LN morphologies to assess the overall impact of synapse placement and input location on dendritic integration.

### Statistical analysis

Statistical analyses were performed using Navis (Python), Natverve (R), and TREES Toolbox (Matlab). We used Pearson’s correlation and Spearman rank correlation to analyse the relationships between various parameters in our datasets. Permutation tests were used to quantify significance levels. To test if two samples came from the same distribution two-sample Kolmogorov–Smirnov (KS) test were used. The number of observations, significance values, and other relevant information for data comparisons are specified in the respective figure legend and in text. In all figures, * represents p value < 0.05, ** represents p value < 0.01, and *** represents p value < 0.001.

## Supplementary figures

**Figure S1.**
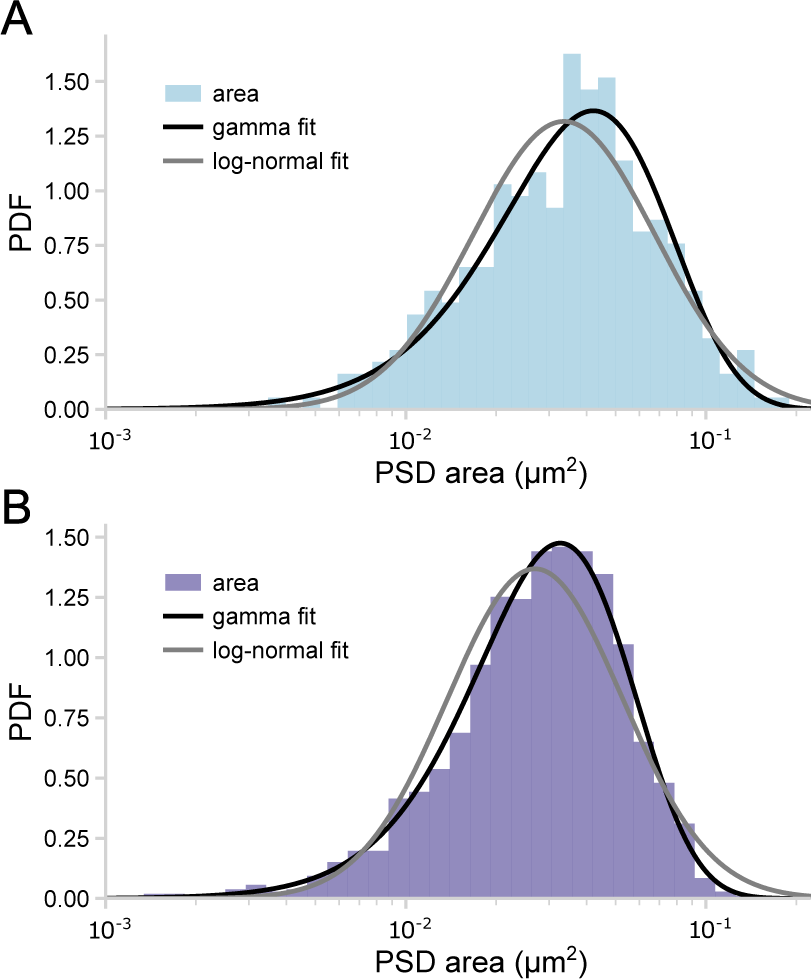
Supplementary to *Figure 2*: PSD area distributions across development with fitted log-normal and gamma models. (**A**,**B**) Synaptic size (PSD area) distributions of L1 (**A**) and L3 (**B**) datasets, both fitted by a gamma distribution (black) and a log-normal distribution (gray).

**Figure S2.**
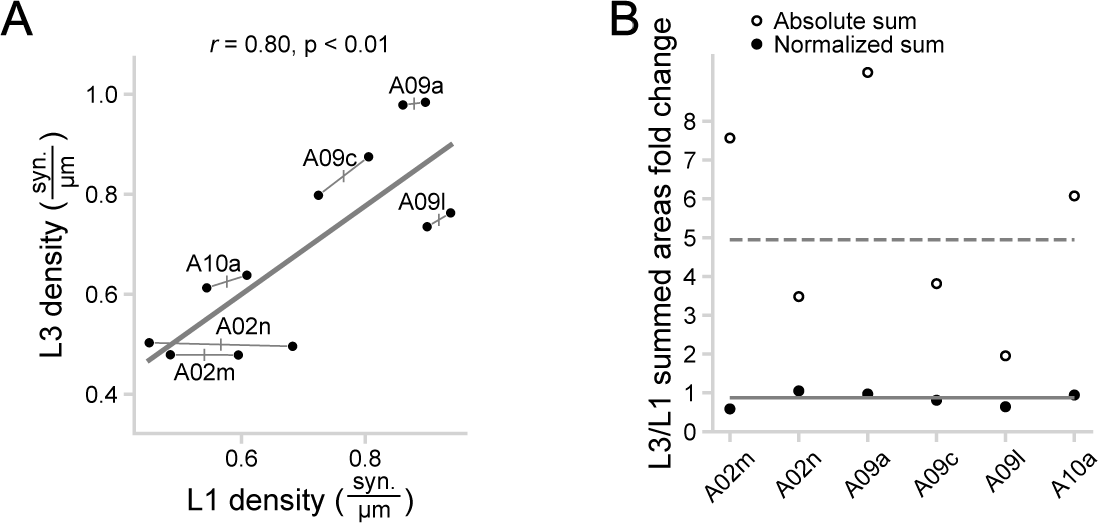
Supplementary to *Figure 4*: Synaptic density conservation across development. (**A**) Scatter plot comparing synaptic density (synapses per *μm*) in L1 and L3 across different LN subtypes. Each LN type is represented by two data points, corresponding to the left and right counterparts of the VNC. Synaptic density is strongly correlated between developmental stages (*r* = 0.80, *p* < 0.01), suggesting that dendritic synaptic density is maintained despite neuronal growth. The solid line represents the linear regression fit. (**B**) Mean fold change (L3/L1) of summed PSD areas from each mdIV–LN connection for different LN subtypes (open circles), along with the fold change of summed PSD areas normalized by the number of synapses within each connection for the same subtypes. The dashed line shows the median 4.95 fold increase in summed PSD areas and grey line shows the median 0.88 fold change of the normalized summed areas. Normalized values remain close to 1, suggesting that the synaptic sizes per connection are stable across development.

**Figure S3.**
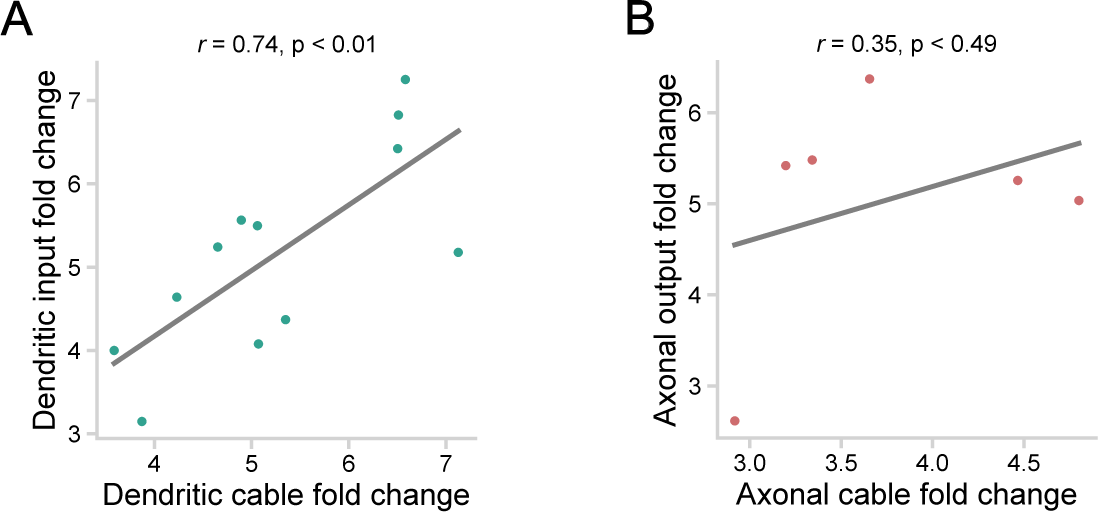
Supplementary to *Figure 5*: Dendritic synaptic input increases proportionally with dendritic growth, while axonal output increases independently of axonal growth. (**A**) Scatter plot showing the relationship between dendritic cable growth ratio (L3/L1) and number of synaptic inputs ratio (L3/L1) across LN subtypes. Input ratio scales proportionally with dendritic growth (*r* = 0.74, *p* < 0.01), suggesting that synaptic input is adjusted to match dendritic expansion. (**B**) Scatter plot of axonal cable growth ratio (L3/L1) versus number of synaptic outputs ratio (L3/L1) for mdIV neurons. Axonal output ratio does not significantly correlate with axonal growth (*r* = 0.35, *p* = 0.49). Solid lines represent linear regression fits.

**Figure S4.**
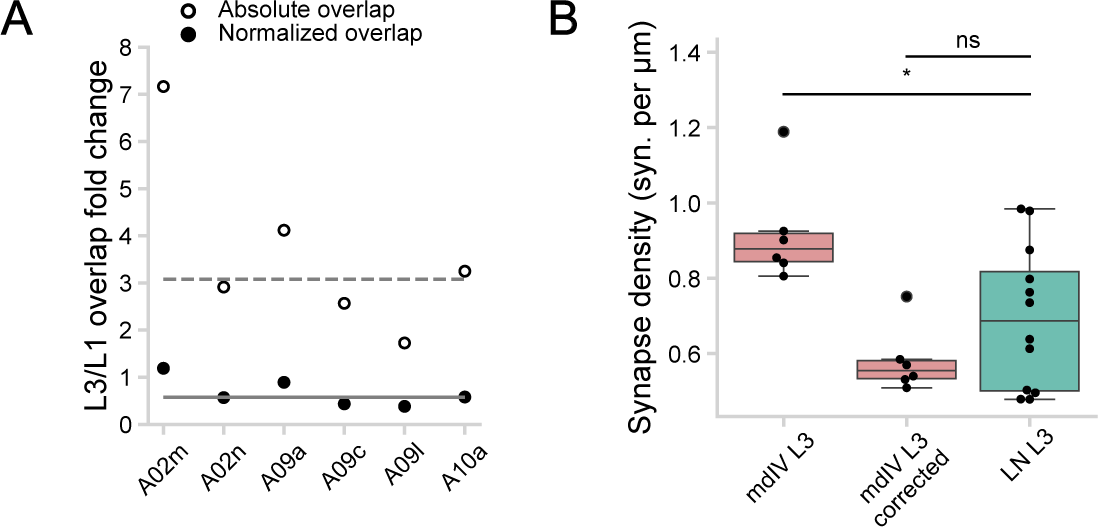
Supplementary to *Figure 6*: Cable overlap and synaptic density comparisons between L1 and L3. (**A**) Fold change in cable overlap (L3/L1) for different LN subtypes. Open circles show absolute cable overlap change, and filled circles show normalized cable overlap. Dashed and gray lines indicate the median fold change in absolute (3.36) and normalized (0.63) overlap, respectively. (**B**) Comparison of synaptic density between presynaptic mdIV axons and postsynaptic LN dendrites in L3. Mean mdIV density (left, red) is significantly higher than dendritic density (right, green; *p* < 0.05). After correcting mdIV density by the median fold change in normalized cable overlap (0.632; middle, green), adjusted densities are not significantly different from dendritic densities (*p* = 0.86).

**Figure S5.**
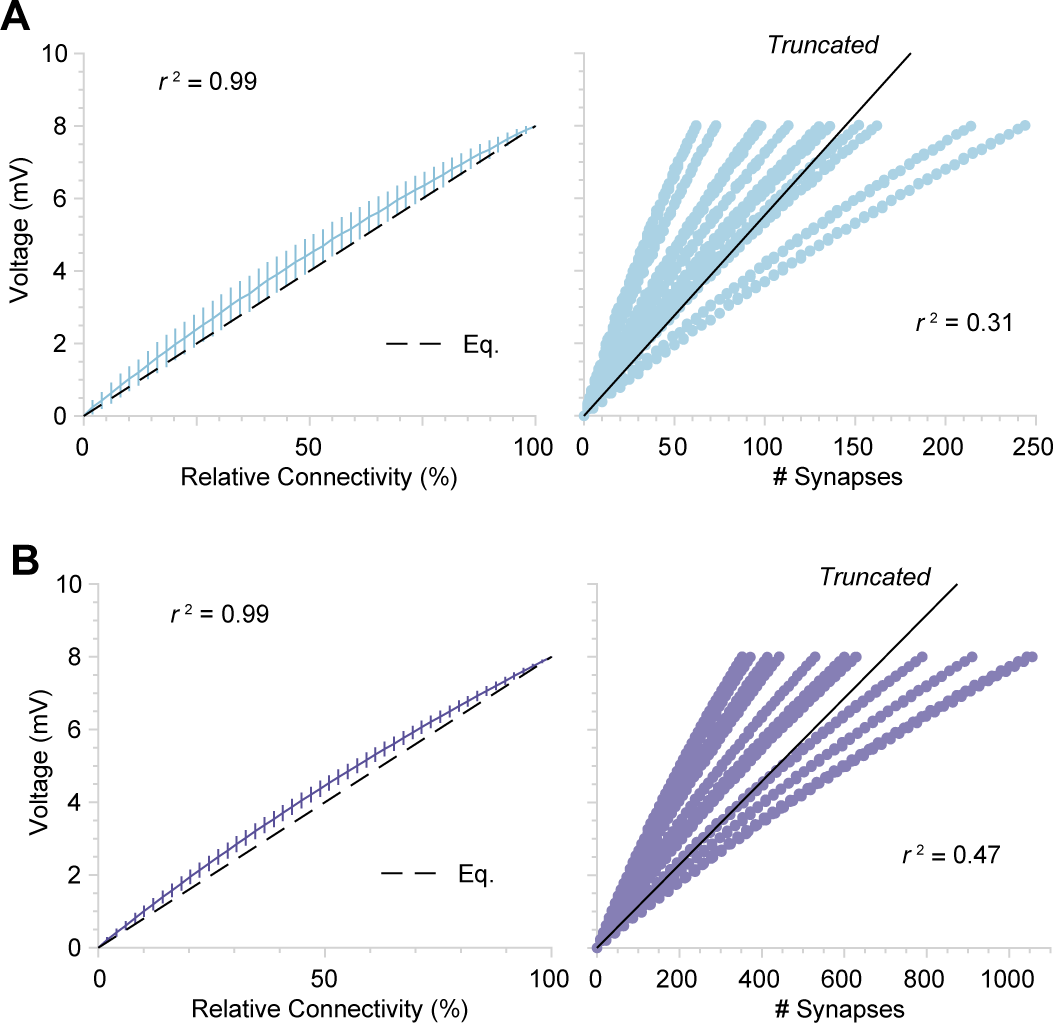
Supplementary to *Figure 7*: Simulated voltage responses as a function of relative connectivity or absolute number of synapses in mdIV-LN connection. (**A**) Simulated steady-statSimulated steady-state voltage responses in each L1 LN neuron. Left: voltage responses across L1 LN morphologies plotted against relative connectivity (percentage of total synaptic input in 2% intervals). The dashed line shows the analytical prediction from the equation (Eq.) 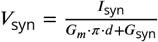, with close agreement between data and prediction (*r*^2^ = 0.99). Right: same data plotted against the absolute number of synapses per LN (*r*^2^ = 0.31). The solid line shows the best-fit truncated regression. (**B**) Same analysis as in (**A**) but for L3 LN morphologies.

